# ALKBH5 Promotes Hepatic Gluconeogenesis via m^6^A-mediated Stabilization of *Ogt* mRNA

**DOI:** 10.64898/2026.03.02.708968

**Authors:** Yan Chen, Yujie Fu, Nan Wei, Xianghong Xie, Yuesi Xu, Wenqi Pan, Qiao Wu, Yifan Li, Qing Li, An lv, Chunying Liu, Hengbo Wu, Hong Liu, Chunpu Li, Wei-Min Tong, Xiaojun Liu, Yamei Niu

## Abstract

Excessive hepatic glucose production is a hallmark of fasting hyperglycemia in type 2 diabetes mellitus, yet the epitranscriptomic mechanisms that sustain this dysregulation remain incompletely understood. Here, we show that dexamethasone/forskolin-induced gluconeogenic activation triggers widespread remodeling of RNA m^6^A methylation in mouse primary hepatocytes. Among the m^6^A regulators examined, the demethylase ALKBH5 is robustly induced both in vitro and in fasted livers. This induction is mediated by the glucocorticoid receptor, which binds to the *Alkbh5* promoter and activates its transcription. Functionally, ALKBH5 overexpression enhances gluconeogenic gene expression and glucose production, whereas both global and hepatocyte-specific *Alkbh5* knockout mice exhibit suppressed hepatic gluconeogenesis. Mechanistically, we identify *Ogt* as a primary downstream target, defining an ALKBH5–OGT epitranscriptomic axis in hepatocytes. ALKBH5 demethylates and stabilizes *Ogt* mRNA, thereby elevating OGT expression and promoting gluconeogenic gene expression. Consistently, the pro-gluconeogenic effects of ALKBH5 are largely abrogated by OGT knockdown. Finally, pharmacological inhibition of ALKBH5 with 18l effectively improves glucose tolerance and suppresses hepatic gluconeogenesis in both wild-type and *db/db* mice. Together, these findings reveal a glucocorticoid-responsive epitranscriptomic mechanism whereby ALKBH5 stabilizes *Ogt* mRNA to drive gluconeogenesis and highlight ALKBH5 as a promising therapeutic target for type 2 diabetes mellitus.

## 1. Introduction

Type 2 diabetes mellitus (T2DM) is characterized by chronic hyperglycemia, driven largely by excessive hepatic glucose production (HGP), which accounts for up to 90% of endogenous glucose production [1, 2]. Among the pathways contributing to HGP, unrestrained activation of hepatic gluconeogenesis represents the primary driver of fasting hyperglycemia [1, 3–6]. Accordingly, suppression of excessive gluconeogenesis remains a cornerstone of current T2DM therapies; however, therapeutic strategies that directly and selectively target this pathway remain limited [7, 8]. Identifying regulatory mechanisms that fine-tune gluconeogenic output is therefore of central importance for improving metabolic intervention.

Given its central contribution to fasting hyperglycemia, hepatic gluconeogenesis must be precisely controlled to adapt glucose output to systemic metabolic demands. This process is tightly controlled by hormonal and nutritional cues, including glucocorticoids and glucagon, acting through transcriptional and post-transcriptional mechanisms to regulate key gluconeogenic enzymes [2, 9]. While transcriptional control of gluconeogenic genes has been extensively studied, post-transcriptional regulatory layers that translate hormonal signals into sustained gluconeogenic flux remain comparatively underexplored, particularly under diabetic conditions.

N^6^-methyladenosine (m^6^A) is the most abundant and conserved internal modification in mammalian mRNAs [10], and serves as a dynamic regulator of RNA splicing, translation, and stability through coordinated action of methyltransferases, demethylases, and m^6^A-binding proteins [11, 12]. Emerging evidence has linked m^6^A modification to metabolic disorders such as T2DM [13], pancreatic β-cell function [14–17], and insulin sensitivity [18–20]. Notably, changes in global m^6^A content and dysregulated expression of m^6^A regulators have been observed in T2DM patients [18, 21, 22]. However, most existing studies rely on bulk measurements of m^6^A levels in patient samples or whole tissues and provided limited insight into the dynamic regulation and functional engagement of the m^6^A machinery during specific metabolic programs. In particular, how m^6^A remodeling is coordinated with gluconeogenic activation in hepatocytes, and how endocrine cues interface with RNA-level regulation of hepatic glucose output, remain incompletely understood.

In this study, we present the m^6^A remodeling landscape during gluconeogenic activation and identify ALKBH5 as a hormone-responsive regulatory factor of hepatic gluconeogenesis. We show that glucocorticoid signaling engages ALKBH5 to modulate hepatic glucose output through m^6^A-dependent post-transcriptional control of metabolic effectors. Together, these findings define a regulatory axis linking endocrine signaling to epitranscriptomic remodeling of hepatic gluconeogenesis and highlight m^6^A regulation as a mechanistic layer governing hepatic glucose metabolism.

## 2. Methods

### 2.1. Animals

All mice had a C57BL/6J genetic background and were maintained in a specific pathogen-free facility under a 12:12 h light/dark cycle with free access to food and water. *Alkbh5* knockout (KO) mice were generated as previously described [23]. Hepatocyte-specific *Alkbh5* knockout (cKO) mice were obtained by crossing *Alkbh5^flox/flox^* mice with *Alb*-*Cre* transgenic mice. For fasting experiments, 8-week-old male WT C57BL/6J mice were deprived of food but allowed free access to water for 48 h. 8-week-old male *db/db* mice were purchased from Baidelong Biotechnology (Hebei, China). For the *db/db* mice experiments, only male animals exhibiting stable hyperglycemia (fasting blood glucose > 250 mg/dL) were included in the study. All experimental animals were acclimated for at least three days prior to the start of any intervention. All animal experiments were performed according to the guidelines of Animal Care and Use Committee of IBMS/CAMS. Ethical approval for the study was obtained from the Institutional Review Board of IBMS/CAMS (Approval Number: ACUC-XMSB-2024-113). The genotypes were determined via PCR using the primers listed in Supplementary Table 1.

### 2.2. Isolation of mouse primary hepatocytes

MPH were isolated from 8-week-old male C57BL/6 or *Alkbh5^flox/flox^* mice as previously described [24], with modifications. Briefly, mice were anesthetized, and the livers were perfused through the inferior vena cava using a disposable indwelling needle, with the portal vein severed to allow drainage. A prewarmed (42 °C) wash buffer (0.34 g/L KCl, 0.18 g/L NaH_2_PO_4_, 7 g/L NaCl, 1.26 g/L NaHCO_3_, 0.1 g/L Na_2_HPO_4_, 1.8 g/L glucose, 0.0468 g/L MgCl_2_, and 0.5 mM EGTA) was infused at 5 mL/min, followed by collagenase buffer (42 °C) containing the same basal salts supplemented with 2.747 mM CaCl_2_ and 100 CDU/mL collagenase type II (Sigma-Aldrich, C6885, USA) for approximately 5 min. The digested livers were excised, gently minced, and filtered through a 70-μm cell strainer (Corning, 352350, USA). Cell suspensions were centrifuged in culture medium at 50 × g for 2 min at 4 °C to remove debris, and the resulting pellets were further purified by centrifugation in 40% Percoll (Cytiva, 17089101, USA) at 200 × g for 5 min. The enriched hepatocytes were seeded into collagen (Corning, 354236) coated plates and cultured in RPMI 1640 medium (National Biomedical Cell Resource Bank (NBCRB), A100009, Beijing, China) containing 10% fetal bovine serum (FBS, AusGeneX, FBS500-S, Australia) and 1% penicillin-streptomycin (Solarbio, P1400, Beijing, China) at 37 °C in a humidified incubator with 5% CO_2_.

### 2.3. Cell culture and treatments

The AML12 cell line (RRID: CVCL_0140) was obtained from the Cell Bank of the Chinese Academy of Sciences / National Experimental Cell Resource Sharing Platform (Catalog No. GNM42, CSTR:19375.09.3101MOUGNM42) in October 2024. The cell line was authenticated by short tandem repeat (STR) profiling provided by the supplier and confirmed to be of murine origin without cross-contamination from other species. Mycoplasma contamination was routinely tested in our laboratory and the cells were confirmed to be free of contamination. AML12 cells were cultured in DMEM/F12 medium (Pricella, PM150312, Wuhan, China) supplemented with 10% FBS, 1% penicillin-streptomycin, 1× insulin-transferrin-selenium (Pricella, PB180429, Wuhan, China) and 40 ng/mL dexamethasone (AbMole, M2176, USA). Cells were maintained at 37 °C in a humidified incubator with 5% CO_2_.

For drug treatment experiments, MPH and AML12 cells were seeded into 12-well culture plates at an appropriate density to achieve ∼70–80% confluency, followed by addition of 1 µM dexamethasone and 10 µM of forskolin (Abmole, M2191, USA) dissolved in DMSO (Solarbio, D8371, Beijing, China). Where necessary, cells were treated with 10 µM mifepristone (Abmole, M3510, USA) dissolved in DMSO for 24 h. Drugs were diluted with RPMI-1640 or DMEM/F12 supplemented with 2% FBS, depending on the cell type. Following treatment, cells were collected either by trypsinization for protein extraction or by direct lysis in TRIzol reagent (Invitrogen, 15596018CN, USA) for RNA isolation.

### 2.4. Other methods

The following methods are described in the Supporting Information, including gene knockdown, knockout and overexpression, RNA extraction and qPCR, m^6^A-sequencing analysis (m^6^A-seq) and m^6^A-IP-qPCR, protein extraction and Western blotting analysis, immunohistochemistry analysis, ChIP assay, dual-luciferase reporter assay, glucose output assay, pyruvate tolerance test (PTT) and glucose tolerance test (GTT), RIP-seq and RIP-qPCR, subcellular fractionation analysis of RNA, RNA stability assay, polysome profiling analysis, cell viability assay, in vitro and in vivo application of 18l, and sequencing data analyses.

### 2.5. Statistical analysis

Data are presented as means ± SEMs. The number of experimental repeats is detailed in the figure legends. Statistical analyses were conducted via GraphPad Prism 10.4.1. Comparisons were performed via two-tailed unpaired Student’s t-test, one-way ANOVA or two-way ANOVA, as specified in the figure legends. A *p*-value of less than 0.05 was considered statistically significant.

## 3. Results

### 3.1. Fasting induces RNA m^6^A dysregulation and ALKBH5 upregulation in mouse primary hepatocytes

To assess m^6^A dynamics during hepatic gluconeogenesis, mouse primary hepatocytes (MPH) were treated with dexamethasone (Dex, a synthetic glucocorticoid receptor agonist) and forskolin (Fsk, an adenylate cyclase activator) to pharmacologically induce gluconeogenesis, followed by m^6^A-seq (Supplementary Fig. 1**a**–**c**, Supplementary Table 2). Comparative methylation analysis revealed a global shift toward reduced m^6^A methylation upon treatment (Fig. 1**a**, Supplementary Fig. 1**d**), indicating disruption of m^6^A homeostasis during gluconeogenic activation. Notably, differential methylation analysis showed that the number of hypomethylated peaks (2 125) markedly exceeded that of hypermethylated ones (650) (Fig. 1**b**). Gene Ontology (GO) analysis showed that, although genes corresponding to hypermethylated or hypomethylated RNAs were largely distinct, both sets were functionally linked to metabolic and stress-responsive pathways, including RNA splicing, autophagy and glycolipid metabolism (Fig. 1**c**, Supplementary Table 3).

**Fig. 1.**
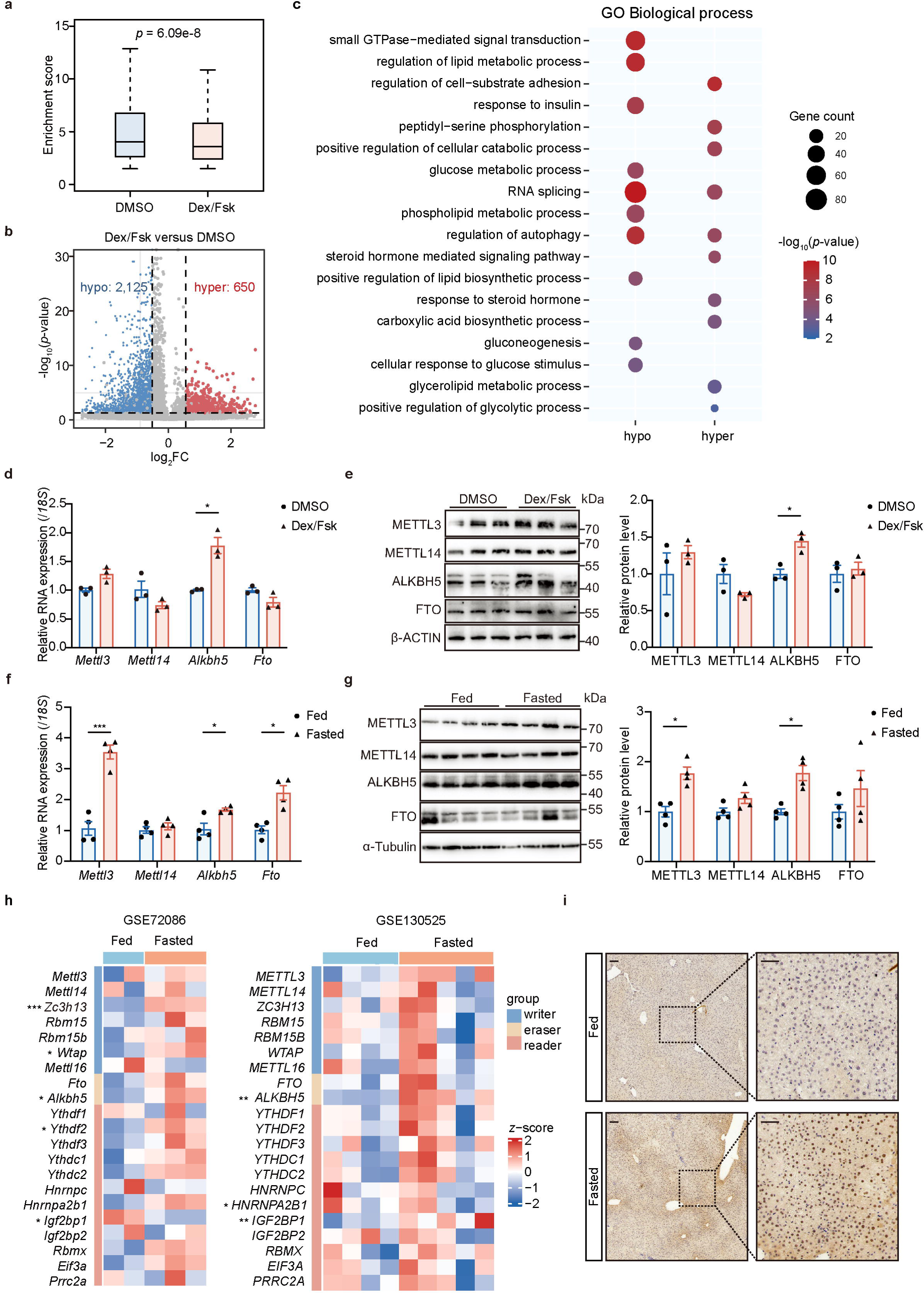
Fasting induces RNA m^6^A dysregulation and ALKBH5 upregulation in mouse primary hepatocytes. **a** Boxplots showing the relative methylation levels of all m^6^A peaks between DMSO- and Dex/Fsk-treated mouse primary hepatocytes (MPH). **b** Volcano plot showing the distribution of differential m^6^A peaks between DMSO- and Dex/Fsk-treated MPH. The black dotted lines indicate a fold change of 1.3 and *p* < 0.05. **c** GO analysis of genes corresponding to hypomethylated or hypermethylated RNAs. **d** RT-qPCR analysis of four key m^6^A regulators in MPH treated with either DMSO or Dex/Fsk (n = 3). **e** Representative Western blot (left) and quantification (right) of four m^6^A regulators in MPH treated with either DMSO or Dex/Fsk (n = 3). **f** RT-qPCR analysis of four key m^6^A regulators in liver tissues from the mice that were fed or fasted for 48 h (n = 4). **g** Representative Western blot (left) and quantification (right) of the four m^6^A regulators in the liver tissues from the mice that were fed or fasted for 48 h (n = 4). **h** Heatmap illustrating the RNA expression levels of m^6^A regulators in MPH from fed and fasted mice in two datasets. GSE72086 contains mouse-derived cells, whereas GSE130525 contains human xenograft-derived cells. **i** Representative images of immunohistochemical staining of ALKBH5 in liver tissues from the mice that were fed or fasted for 48 h (n = 4). Scale bars: 100 μm (left panel), 10 μm (right panel). Data are represented as mean ± SEM. Statistical significance was calculated by two-tailed unpaired Student’s t-test (**a**) or multiple unpaired Student’s t-tests with Holm-Sidak correction (**d**–**g**). Differential expression: Wald test (DESeq2, GSE130525) or t-test (log_2_FPKM, GSE72086), both with Benjamini-Hochberg correction (**h**). \**p* < 0.05, \*\**p* < 0.01, and \*\*\**p* < 0.001.

To investigate the mechanism underlying m^6^A dysregulation during hepatic gluconeogenesis, we examined the expression of m^6^A regulators in response to Dex/Fsk treatment. Among the four key regulators examined here (METTL3, METTL14, ALKBH5 and FTO), only ALKBH5 exhibited consistent induction at both the mRNA (Fig. 1**d**, Supplementary Fig. 1**e**) and protein levels (Fig. 1**e**). In the livers of fasted mice, ALKBH5 expression was similarly upregulated (Fig. 1**f**–**g**). Furthermore, re-analysis of previously published RNA-seq data of livers from fasted mice (GSE72086) and a humanized liver mouse model (GSE130525) confirmed selective induction of ALKBH5 under fasting conditions (Fig. 1**h**) [25, 26]. Consistently, immunohistochemical analysis confirmed an increase in ALKBH5 expression in liver sections from fasted mice (Fig. 1**i**).

Taken together, these results suggest that gluconeogenic activation is associated with global alterations in RNA m^6^A methylation, potentially mediated by the upregulation of the demethylase ALKBH5 in hepatocytes.

### 3.2. Glucocorticoid receptor enhances *Alkbh5* gene transcription

Since Dex/Fsk treatment increased ALKBH5 expression, we next sought to investigate the mechanism underlying ALKBH5 upregulation. Dex, but not Fsk, significantly elevated *Alkbh5* mRNA expression in MPH (Fig. 2**a**), which was further validated in another mouse hepatocyte cell line, AML12 (Fig. 2**b**). Given that Dex activates GR signaling [27], we further investigated whether GR directly regulates *Alkbh5* transcription. GR overexpression increased the mRNA levels of *Alkbh5* in both MPH and AML12 cells (Supplementary Fig. 2**a**–**c**, Fig. 2**c**–**d**). Conversely, treatment with the GR inhibitor mifepristone markedly reduced *Alkbh5* expression in both cell types (Supplementary Fig. 2**d**, Fig. 2**e**–**f**).

**Fig. 2.**
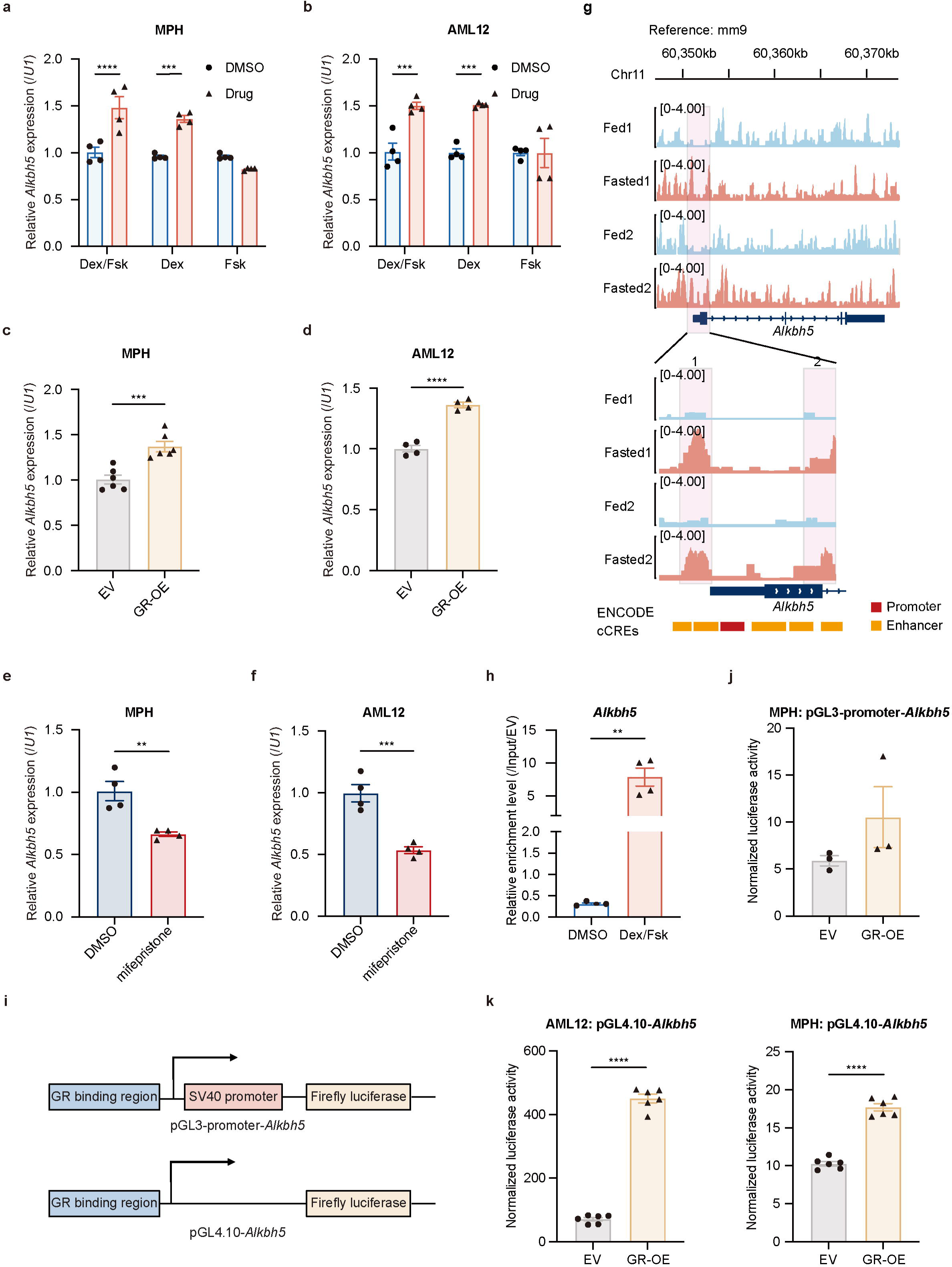
Glucocorticoid receptor enhances the transcription of *Alkbh5* gene. **a**–**b** RT-qPCR analysis of *Alkbh5* expression in MPH (**a**) and AML12 cells (**b**) treated with DMSO, Dex/Fsk, Dex or Fsk (n = 4). **c**–**d** RT-qPCR analysis of *Alkbh5* expression in GR-overexpressing (GR-OE) MPH (**c**) and AML12 cells (**d**) treated with Dex (n = 6, n = 4, respectively). **e**–**f** RT-qPCR analysis of *Alkbh5* expression in GR inhibitor mifepristone-treated MPH (**e)** and AML12 cells (**f**) (n = 4). **g** IGV tracks depicting genome-wide GR ChIP-seq profiles at the *Alkbh5* gene locus in liver tissues from fed and fasted mice (GSE72087), alongside candidate cis-regulatory elements (cCREs) annotated by the ENCODE project. An enlarged view of the shaded region highlights two potential GR-binding regions upstream of the *Alkbh5* gene. **h** ChIP-qPCR of the putative GR-binding region within *Alkbh5* in AML12 cells. Enrichment was calculated relative to input and empty vector (EV) (n = 4). **i** Schematic of luciferase reporter constructs with the GR-binding region of *Alkbh5* cloned into pGL3-promoter and pGL4.10 vectors. **j** Dual-luciferase reporter assay to compare EV and GR-overexpressing (OE) groups by using MPH (n = 3) that were transfected with pGL3-promoter-*Alkbh5* and treated with Dex/Fsk (n = 3). **k** Dual-luciferase reporter assay to compare EV and GR-OE groups by using MPH (n = 6) and AML12 cells (n = 4) that were transfected with pGL4.10-*Alkbh5* and treated with Dex/Fsk. Data are represented as mean ± SEM. Statistical significance was calculated by two-way ANOVA followed by Sidak’s multiple comparisons test (**a**–**b**) or two-tailed unpaired Student’s t-test (**c**–**f**, **h**, **j**–**k**). \**p* < 0.05, \*\**p* < 0.01, and \*\*\**p* < 0.001.

Re-analyzed previously published GR- chromatin immunoprecipitation (ChIP)-seq datasets from the mouse liver tissues (GSE72087 and GSE137978) [25, 28] identified multiple GR-binding regions within the *Alkbh5* locus. Notably, under fasting conditions, GR binding was significantly enhanced at two upstream regions (Fig. 2**g**). Integrating analysis with assay for transposase-accessible chromatin (ATAC) sequencing data of mouse liver from the ENCODE3 project [29–31] showed that only the first of these two regions displayed increased chromatin accessibility (Supplementary Fig. 2**e**). Consistently, our ChIP-qPCR analysis confirmed robust GR enrichment at this region following Dex/Fsk treatment (Fig. 2**h**, Supplementary Fig. 2**f**). According to ENCODE functional annotations [32], this GR-binding region may act as a cis-regulatory element, potentially acting as either a promoter or an enhancer of *Alkbh5* expression (Fig. 2**g**, Supplementary Fig. 2**e**). To experimentally determine its regulatory nature, we employed two types of luciferase reporter systems (Fig. 2**i**). In a SV40 promoter-driven luciferase reporter system, insertion of the GR-binding region did not enhance luciferase activity upon GR overexpression, arguing against enhancer activity (Fig. 2**j**). In contrast, insertion of the same GR-binding sequence into a promoter-less luciferase construct, upstream of the luciferase gene conferred strong GR-dependent transcriptional activation, confirming its promoter activity (Fig. 2**k**).

Collectively, these findings establish *Alkbh5* as a direct transcriptional target of glucocorticoid signaling, establishing a direct transcriptional link between hormonal cues and epitranscriptomic remodeling during gluconeogenic activation.

### 3.3. Hepatic ALKBH5 deficiency ameliorates gluconeogenic capacity

In light of ALKBH5 upregulation during hepatic gluconeogenesis, we next investigated its functional significance in this metabolic process. ALKBH5 overexpression in MPH led to increased mRNA and protein levels of key gluconeogenic enzymes PCK1 and G6PC (Fig. 3**a**–**b**), accompanied by enhanced glucose production (Fig. 3**c**). Conversely, genetic ablation of ALKBH5 in MPH (Supplementary Fig. 3**a**) resulted in reduced expression of PCK1 and G6PC, along with decreased glucose production (Fig. 3**d**–**f**).

**Fig. 3.**
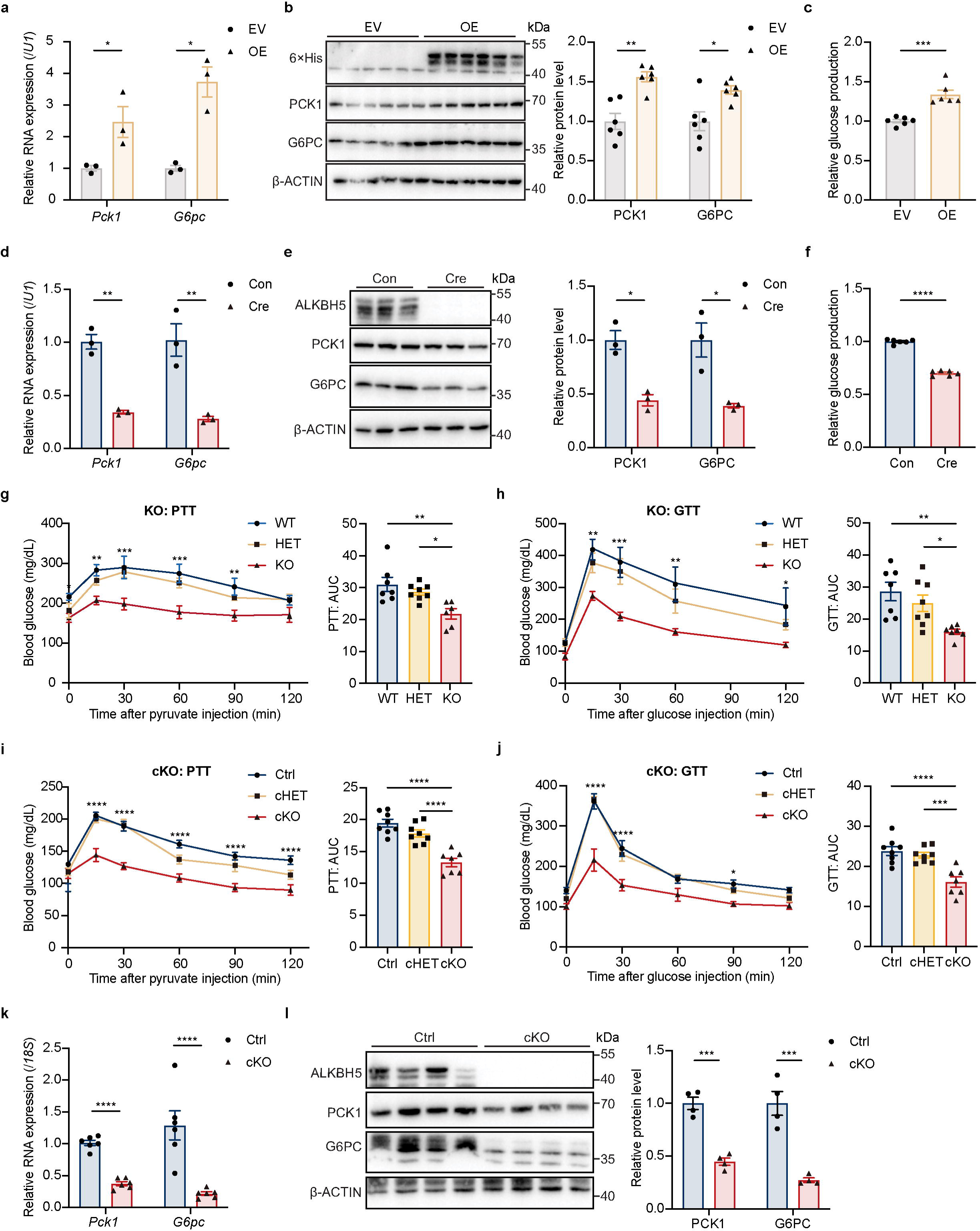
Hepatic ALKBH5 deficiency ameliorates gluconeogenesis. **a** RT-qPCR analysis of *Pck1* and *G6pc* expression in MPH that were transfected with empty vector (EV) or ALKBH5-overexpressing (OE) plasmid (n = 3). **b** Representative Western blot (left) and quantification (right) of PCK1 and G6PC in MPH that were transfected with EV or ALKBH5-OE plasmid (n = 6). **c** Glucose output assay in MPH that were transfected with EV or ALKBH5-OE plasmid (n=6). **d**–**f** RT-qPCR analysis of *Pck1* and *G6pc* expression (**d**, n = 3), representative Western blot (left) and quantification (right) of PCK1 and G6PC (**e**, n = 3) and glucose output assay (**f**, n = 6) in MPH isolated from *Alkbh5^flox/flox^*mice and infected with control (Con) or Cre-expressing (Cre) adenovirus. **g**–**h** Pyruvate tolerance test (PTT) (**g**) and glucose tolerance test (GTT) (**h**) in 6-month-old wild-type (WT), *Alkbh5*-heterozygote (HET) and *Alkbh5*-knockout (KO) mice following a 6-hour fasting. Left: blood glucose concentrations over time. Right: AUC analysis (n: **g**: WT: HET: KO = 7: 8: 6, **h**: WT: HET: KO = 7: 8: 7). **i**–**j** PTT (**i**) and GTT (**j**) in 6-month-old littermate control (Ctrl), hepatocyte-specific *Alkbh5*-heterozygote (cHET), *Alkbh5*-knockout (cKO) mice following a 6-hour fasting (n: Ctrl: cHET: cKO = 8: 8: 7). **k** RT-qPCR analysis of *Pck1* and *G6pc* expression in liver tissues from Ctrl and *Alkbh5*-cKO mice (n = 6). **l** Representative Western blot (left) and quantification (right) of PCK1 and G6PC in liver tissues from Ctrl and *Alkbh5*-cKO mice (n = 4). Data are represented as mean ± SEM. Statistical significance was calculated by multiple unpaired Student’s t-tests with Holm-Sidak correction (**a**–**b**, **d**–**e**, **k**–**l**), two-tailed unpaired Student’s t-test (**c**, **f**), two-way ANOVA followed by Dunnett’s multiple comparisons test (blood glucose concentrations in **g**–**j**) or one-way ANOVA with Dunnett’s multiple comparisons test (AUC in **g**–**j**). \**p* < 0.05, \*\**p* < 0.01, and \*\*\**p* < 0.001.

To evaluate the systemic impact of ALKBH5 on glucose metabolism, we next analyzed global *Alkbh5*-knockout (KO) [23] mice under normal chow diet conditions. Despite similar body weights and liver weights (Supplementary Fig. 3**b**–**d**), *Alkbh5*-deficiency markedly attenuated pyruvate-induced gluconeogenesis (Fig. 3**g**) and improved glucose tolerance (Fig. 3**h**), highlighting its role in systemic glucose homeostasis. To determine whether these metabolic effects are hepatocyte-autonomous, we crossed *Alkbh5^flox/flox^* mice with Albumin-Cre (*Alb*-*Cre*) transgenics to generate hepatocyte-specific *Alkbh5*-knockout mice (*Alkbh5^flox/flox^; Alb-Cre*, cKO) (Supplementary Fig. 3**e**–**f**). Consistent with the global knockout phenotype, hepatocyte-specific deletion of *Alkbh5* also led to decreased pyruvate-induced gluconeogenesis and improved glucose tolerance (Fig. 3**i**–**j**), while the body weights and liver weights remained unaffected (Supplementary Fig. 3**g**–**i**). Notably, hepatic PCK1 and G6PC expression in cKO mice were significantly downregulated compared to their respective controls (Fig. 3**k**–**l**).

Together, these findings demonstrate that ALKBH5 acts as a positive regulator of hepatic gluconeogenic capacity in a hepatocyte-autonomous manner.

### 3.4. ALKBH5 deficiency caused m^6^A dysregulation in mouse primary hepatocytes

Given the robust effect of ALKBH5 in promoting gluconeogenesis, we next performed m^6^A-seq using *Alkbh5*-deficient MPH to explore the underlying mechanism (Supplementary Fig. 4**a**–**c**, Supplementary Table 2). *Alkbh5*-deficient MPH exhibited a larger number of hypermethylated peaks (1,264) compared to controls than hypomethylated ones (401) (Fig. 4**a**). GO analysis showed that many genes corresponding to those hypermethylated RNAs were functionally relevant with glucose metabolism, such as response to insulin and gluconeogenesis. In contrast, hypomethylated RNAs were enriched in pathways with less direct involvement in glucose metabolism, such as RNA transcription and Rho protein signal transduction. (Fig. 4**b**, Supplementary Fig. 4**d**, Supplementary Table 4). To further define the direct targets of ALKBH5, we performed RNA immunoprecipitation (RIP)-seq in WT MPH (Supplementary Fig. 4**a** and Table 2). Notably, these bound RNAs were also enriched in the biological processes related to metabolism, such as response to insulin and glucose homeostasis (Fig. 4**c**, Supplementary Table 5).

**Fig. 4.**
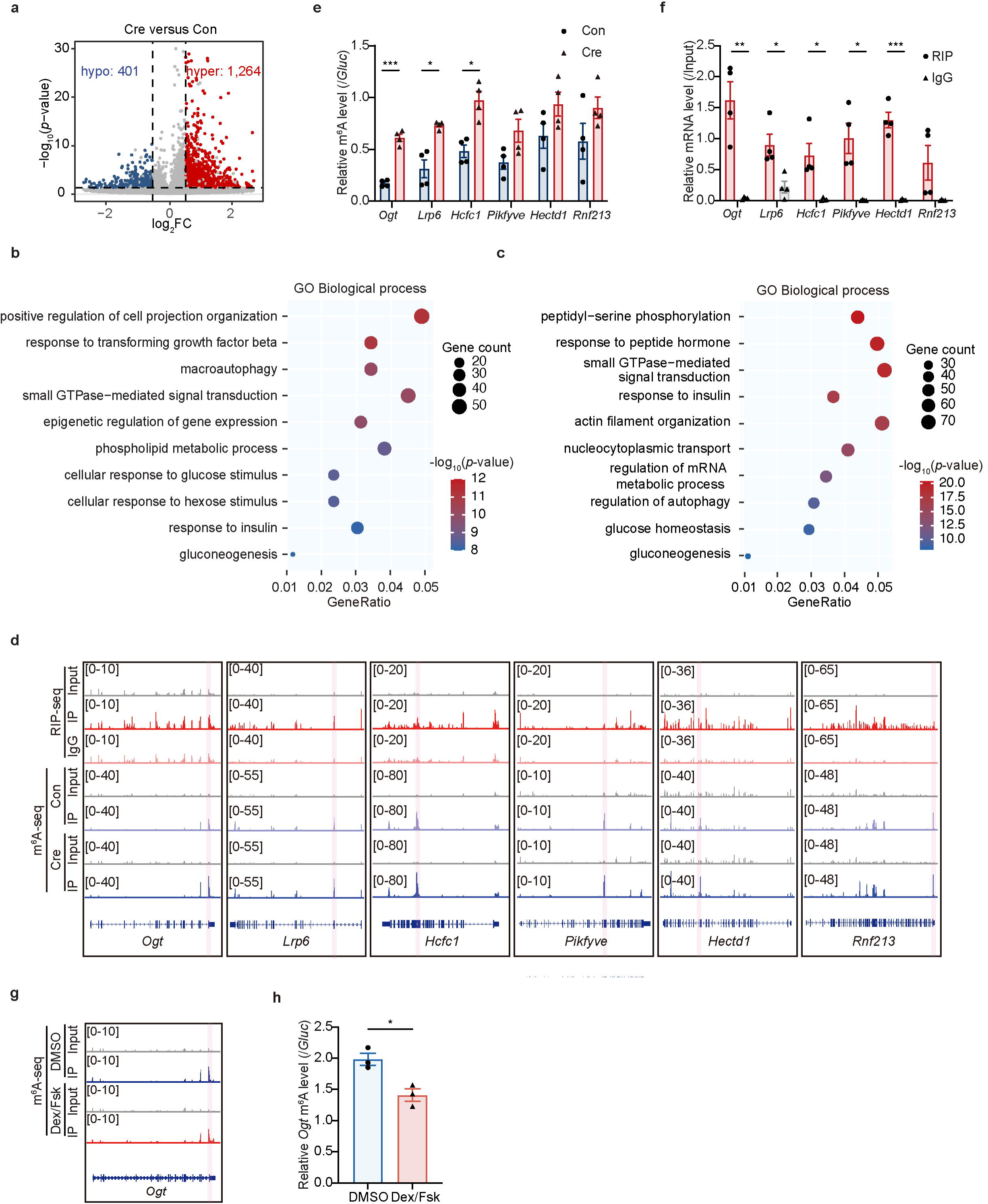
ALKBH5 deficiency causes m^6^A dysregulation in mouse primary hepatocytes. **a** Volcano plot showing the distribution of differential m^6^A peaks between control- (Con) and Cre-expressing (Cre) adenovirus-infected MPH. The black dotted lines indicate a fold change of 1.3 and *p* < 0.05. **b** GO analysis of genes with increased RNA m^6^A methylation in *Alkbh5*-deficient MPH. **c** GO analysis of genes corresponding to ALKBH5-binding RNAs in WT MPH. **d** IGV tracks showing m^6^A profiles and ALKBH5-binding regions in the six potential substrate RNAs of ALKBH5 (*Ogt*, *Lrp6*, *Hcfc1*, *Pikfyve*, *Hectd1*, *Rnf213*). The validated m^6^A peaks are highlighted in red. **e** m^6^A-IP-qPCR results showing the effect of ALKBH5 knockout on the m^6^A levels of indicated transcripts in MPH (n = 4). **f** RIP-qPCR results showing the interaction between ALKBH5 protein and indicated RNAs in WT MPH. Enrichment was calculated relative to input, with IgG as a negative control (n = 4). **g** IGV tracks showing m^6^A profiles of *Ogt* transcript in Dex/Fsk-treated MPH as determined by m^6^A-seq. **h** m^6^A-IP-qPCR results showing the effect of Dex/Fsk treatment on the m^6^A levels of *Ogt* transcript in MPH (n = 3). Data are represented as mean ± SEM. Statistical significance was calculated by multiple unpaired Student’s t-tests with Holm-Sidak correction (**e**–**f**) or two-tailed unpaired Student’s t-test (**h**). \**p* < 0.05, \*\**p* < 0.01, and \*\*\**p* < 0.001.

Among the ALKBH5-bound RNAs identified here, 351 RNAs showed elevated m^6^A levels upon *Alkbh5* deletion (Supplementary Table 6), suggesting that they may represent direct m^6^A demethylating targets of ALKBH5. To narrow down candidate targets for validation, we applied three selection criteria: the magnitude of m^6^A level changes, the strength of ALKBH5-RNA binding signals from RIP-seq, and the known or predicted functions of the encoded proteins. Based on these criteria, we prioritized and selected the top 6 candidate genes—*Ogt*, *Lrp6*, *Hcfc1*, *Pikfyve*, *Hectd1*, and *Rnf213*—for experimental validation using m^6^A-IP-qPCR and RIP-qPCR (Fig. 4**d**). Among the validated candidates, *Ogt* RNA showed marked increase in m^6^A levels upon *Alkbh5* deficiency (Fig. 4**e**), a decrease in m^6^A levels in *Alkbh5*-overexpression MPH (Supplementary Fig. 4**e**), and robust binding with ALKBH5 protein (Fig. 4**f**, Supplementary Fig. 4**f**). Additionally, consistent with *Alkbh5* upregulation, we also observed a reduced m^6^A level in *Ogt* in Dex/Fsk-treated MPH (Fig. 4**g–h**). Taken together with its established role in glucose metabolism [33, 34], *Ogt* RNA was prioritized for subsequent mechanistic analysis.

Collectively, our integrative epitranscriptomic analyses reveal that ALKBH5 broadly modulates m^6^A marks on transcripts associated with glucose metabolism, among which *Ogt* emerges as a direct target, warranting focused mechanistic investigation.

### 3.5. ALKBH5 promotes gluconeogenesis via OGT

Having established *Ogt* as a downstream target of ALKBH5, we next explored how m^6^A demethylation modulates *Ogt* RNA metabolism to impact gluconeogenesis. First, transcriptome-wide analysis showed that although ALKBH5 deficiency altered the alternative splicing of 1,073 RNAs, involving 1,352 events (Supplementary Fig. 5**a**), it did not affect *Ogt* splicing (Supplementary Fig. 5**b**). Next, nuclear-cytoplasmic fractionation analysis showed that *Ogt* RNA subcellular localization remained unchanged (Supplementary Fig. 5**c**). Similarly, polysome profiling analysis showed that ALKBH5 did not affect the translation efficiency of *Ogt* RNA either (Supplementary Fig. 5**d**–**f**). However, RNA stability assay showed that *Ogt* transcripts exhibited a longer half-life in ALKBH5-overexpressing MPH (Fig. 5**a**), suggesting that ALKBH5 enhances *Ogt* RNA stability via m^6^A-dependent degradation. Supporting this, we found that YTHDF2 binds to *Ogt* RNA and mediates its m^6^A-dependent RNA decay (Fig. 5**b**–**c**, Supplementary Fig. 5**g**).

**Fig. 5.**
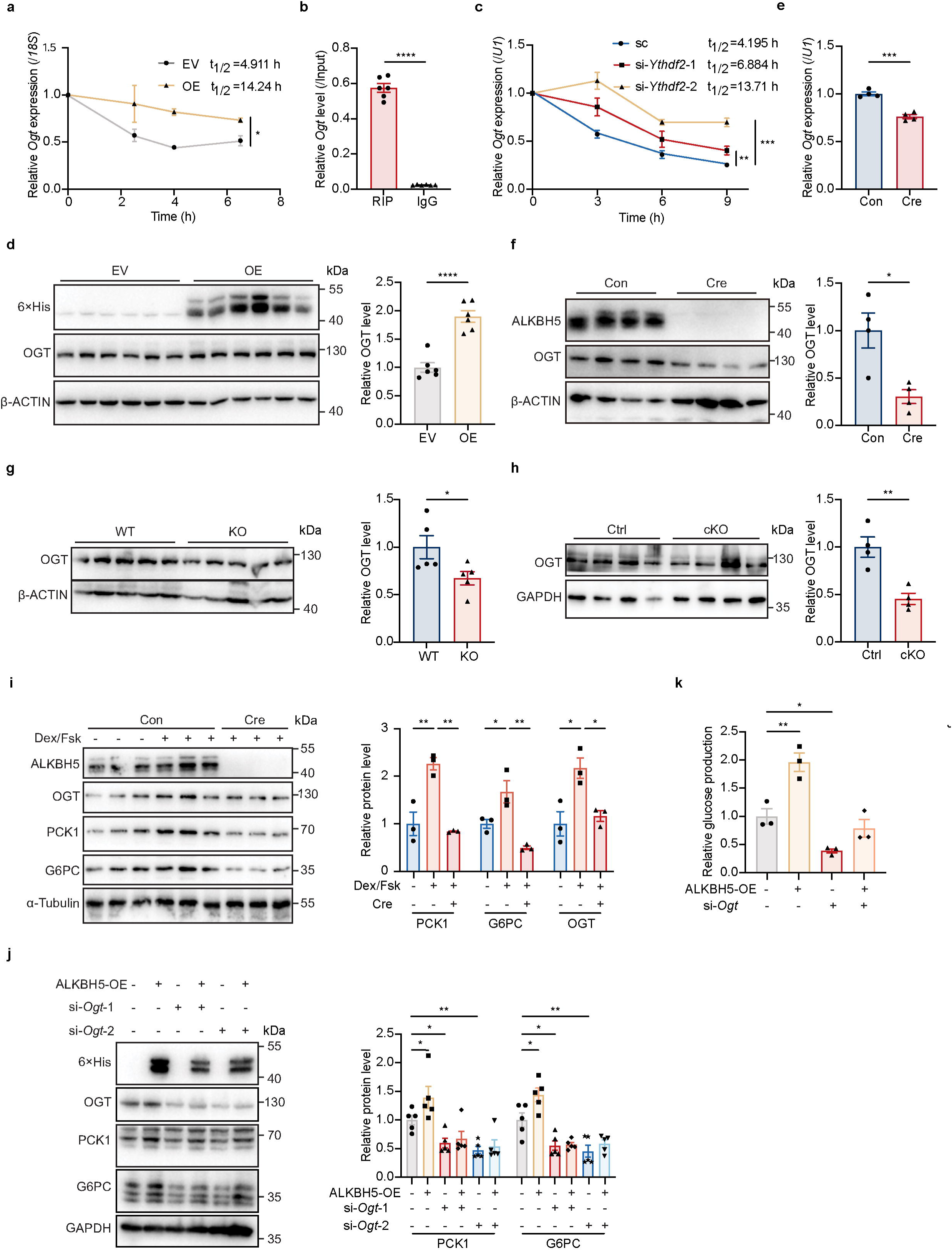
ALKBH5 promotes hepatic gluconeogenesis via OGT. **a** RNA stability assay to assess the effect of ALKBH5 overexpression on the half-life (t_1/2_) of *Ogt* in MPH (n = 4). **b** RIP-qPCR results showing the interaction between YTHDF2 protein and *Ogt* in MPH. Enrichment was calculated relative to input, with IgG as a negative control (n = 6). **c** RNA stability assay to assess the effect of YTHDF2 knockdown on the half-life (t_1/2_) of *Ogt* in MPH (n = 3). **d** Representative Western blot (left) and quantification (right) to detect the effect of ALKBH5 overexpression on OGT expression in MPH (n = 6). **e** RT-qPCR analysis to determine the effect of ALKBH5 knockout on *Ogt* expression in MPH (n = 4). **f** Representative Western blot (left) and quantification (right) to detect the effect of ALKBH5 knockout on OGT expression in MPH (n = 4). **g** Representative Western blot (left) and quantification (right) of OGT in liver tissues from WT and *Alkbh5*-KO mice (n = 5). **h** Representative Western blot (left) and quantification (right) of OGT in liver tissues from Ctrl and *Alkbh5*-cKO mice (n = 4). **i** Representative Western blot (left) and quantification (right) of OGT in *Alkbh5^flox/flox^* MPH treated with DMSO or Dex/Fsk, as well as in Dex/Fsk-treated MPH infected with Cre-expressing adenovirus (n = 3). **j** Representative Western blot (left) and quantification (right) of PCK1 and G6PC in ALKBH5-overexpressing AML12 cells transfected with scramble siRNA or *Ogt* siRNA (n = 5). **k** Glucose output assay in MPH with or without ALKBH5 overexpression, transfected with either scramble siRNA or *Ogt* siRNA (n = 3). Data are represented as mean ± SEM. RNA half-lives were calculated by nonlinear regression (one-phase exponential decay) with group comparisons by extra sum-of-squares F test (**a**, **c**). Other statistical significance was calculated by two-tailed unpaired Student’s t-test (**b**, **d**–**h**), one-way ANOVA with Dunnett’s multiple comparisons test (**i**, **k**) or two-way ANOVA followed by Dunnett’s multiple comparisons test (**j**). \**p* < 0.05, \*\**p* < 0.01, and \*\*\**p* < 0.001.

Consistently, ALKBH5 overexpression increased OGT expression in MPH (Fig. 5**d**, Supplementary Fig. 5**h**), while *Alkbh5* knockout led to a significant reduction in OGT levels (Fig. 5**e**–**f**). Reduced OGT expression was also observed in the livers of both KO and cKO mice (Fig. 5**g**–**h**). Similarly, Dex/Fsk treatment also increased OGT expression in MPH (Fig. 5**i**, Supplementary Fig. 5**i**–**j**), mirroring its effects on ALKBH5, PCK1 and G6PC. However, this effect was alleviated in *Alkbh5*-deficient MPH (Fig. 5**i**).

Functionally, *Ogt* knockdown in MPH decreased G6PC and PCK1 expression and reduced glucose production (Supplementary Fig. 5**k**–**m**), whereas *Ogt* overexpression had the opposite effect (Supplementary Fig. 5**n**–**p**). Importantly, *Ogt* knockdown attenuated the ALKBH5-induced increase in gluconeogenic gene expression and glucose production (Fig. 5**j**–**k**). Together, these findings indicate that OGT acts as a critical downstream effector of ALKBH5 in regulating hepatic gluconeogenesis.

### 3.6. The ALKBH5 inhibitor 18l ameliorates hepatic gluconeogenesis in wild-type and *db/db* mice

Given the established role of ALKBH5 in promoting hepatic gluconeogenesis, we investigated whether its pharmacological inhibition could ameliorate glucose metabolism in T2DM. Building on our previous discovery of 18l as a novel ALKBH5 inhibitor with antileukemia activity [35], we further evaluated its efficacy in hepatocytes in this study. In both MPH and AML12 hepatocytes, cell viability was consistently above 80% throughout 0 to 10 μM (Supplementary Fig. 6**a**–**b**). Notably, treatment with 18l significantly suppressed glucose production and downregulated the expression of G6PC, PCK1 and OGT in hepatocytes (Fig. 6**a**–**c**).

**Fig. 6.**
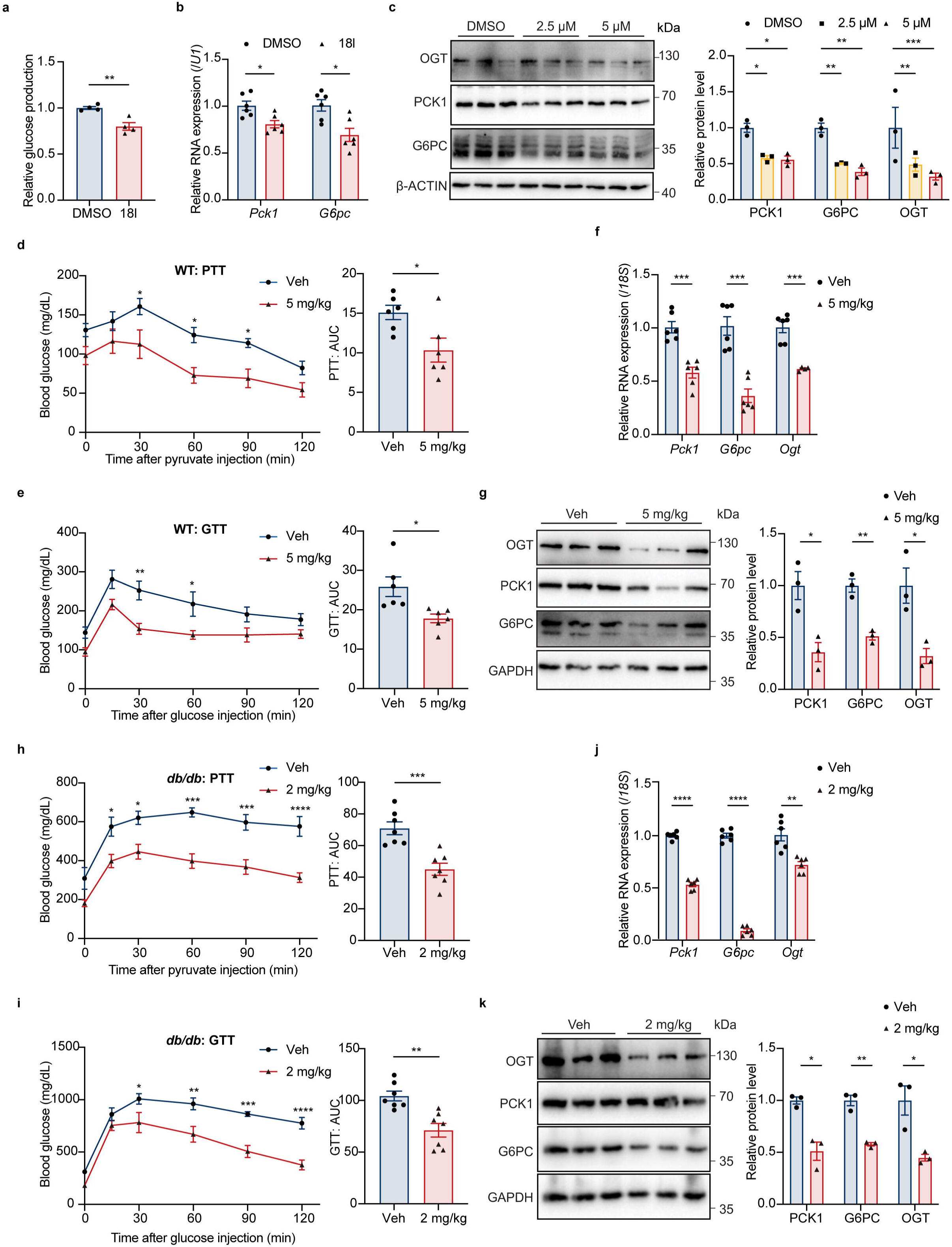
The ALKBH5 inhibitor 18l ameliorates hepatic gluconeogenesis in WT and *db/db*mice. **a** Glucose output assay in MPH treated with 5 μM 18l for 48 h (n = 5). **b** RT-qPCR analysis of *Pck1* and *G6pc* expression in MPH treated with 10 μM 18l (n = 6). **c** Representative Western blot (left) and quantification (right) of PCK1, G6PC and OGT in DMSO, 2.5 μM or 5 μM 18l-treated AML12 cells (n = 3). **d**–**e** PTT and GTT in 2-month-old WT mice, with or without 5 mg/kg 18l injection following 6 h of fasting (n = 6). Left: blood glucose concentrations over time. Right: AUC analysis. **f** RT-qPCR analysis of *Pck1*, *G6pc* and *Ogt* expression in liver tissues from WT mice injected with 5 mg/kg 18l or vehicle (n = 4). **g** Representative Western blot (left) and quantification (right) of PCK1, G6PC and OGT in liver tissues from WT mice injected with 5 mg/kg 18l or vehicle (n = 3). **h**–**i** PTT and GTT in 2-month-old *db/db* mice, with or without 2 mg/kg 18l injection following 16 h of fasting (n = 7). **j** RT-qPCR analysis of *Pck1*, *G6pc* and *Ogt* expression in liver tissues from *db/db* mice injected with 2 mg/kg 18l or vehicle (n = 6). **k** Representative Western blot (left) and quantification (right) of PCK1, G6PC and OGT in liver tissues from *db/db* mice injected with 2 mg/kg 18l or vehicle (n = 3). Data are represented as mean ± SEM. Statistical significance was calculated by two-tailed unpaired Student’s t-test (**a**, AUC in **d**–**e**, **h**–**i**), multiple unpaired Student’s t-tests with Holm-Sidak correction (**b**, **f**–**g**, **j**–**k**), two-way ANOVA followed by Dunnett’s multiple comparisons test (c) or two-way ANOVA followed by Sidak’s multiple comparisons test (blood glucose concentrations in **d**–**e**, **h**–**i**). \**p* < 0.05, \*\**p* < 0.01, and \*\*\**p* < 0.001.

Next, to extend our in vitro findings to an in vivo setting, we evaluated the effects of 181 at 5 mg/kg on hepatic glucose homeostasis and safety in WT mice, with the dosage determined based on our previous leukemia xenograft studies [35]. This treatment significantly attenuated systemic gluconeogenesis while improving glucose tolerance (Fig. 6**d**–**e**). Consistently, hepatic expression of PCK1, G6PC, and OGT was also markedly reduced following treatment (Fig. 6**f**–**g**). No signs of toxicity were observed, as evidenced by unchanged body weights, normal liver function and blood biochemistry, as well as the absence of histopathological abnormalities in major organs (Supplementary Fig. 6**c**–**f**).

Subsequently, for therapeutic evaluation in *db*/*db* mice, we chose a lower dose (2 mg/kg) to minimize potential long-term toxicity and ensure safety during repeated administration. Remarkably, administration of 2 mg/kg of 18l effectively suppressed systemic gluconeogenesis and improved glucose tolerance in *db/db* mice (Fig. 6**h**–**i**), with no obvious change in neither body nor liver weights (Supplementary Fig. 6**g**). Mechanistically, this was accompanied by downregulation of hepatic PCK1, G6PC, and OGT (Fig. 6**j**–**k**).

Together, these results demonstrate that pharmacological inhibition of ALKBH5 by 18l effectively suppresses hepatic gluconeogenesis and improves systemic glucose homeostasis, supporting the translational relevance of targeting ALKBH5-mediated epitranscriptomic regulation in hepatic gluconeogenesis T2DM.

## 4. Discussion

In this study, we show that gluconeogenic stimulation is accompanied by transcriptome-wide RNA m^6^A remodeling in hepatocytes, driven in part by GR-dependent upregulation of ALKBH5. Mechanistically, we identify the ALKBH5–OGT axis as a direct epitranscriptomic link between hormonal cues and hepatic glucose metabolism. We show that ALKBH5 promotes hepatic gluconeogenesis by demethylating and stabilizing *Ogt* mRNA, thereby increasing OGT abundance and enhancing downstream gluconeogenic genes. Functionally, genetic ablation of ALKBH5 suppresses hepatic gluconeogenesis, and pharmacological inhibition of ALKBH5 ameliorates gluconeogenesis in diabetic mouse models, underscoring the translational potential of modulating this pathway in T2DM (Fig. 7).

**Fig. 7.**
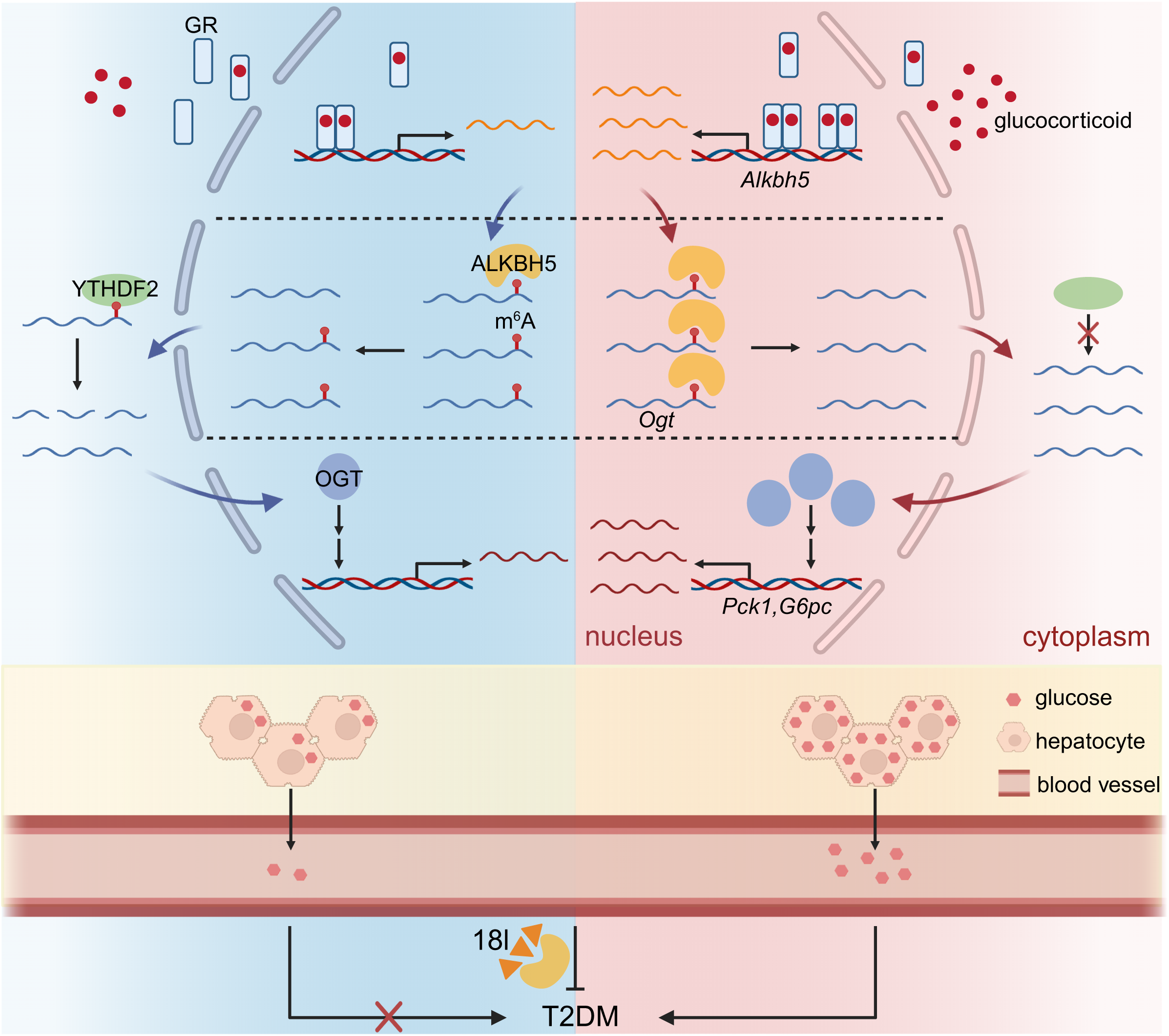
Schematic model of ALKBH5-mediated regulation of hepatic gluconeogenesis via m^6^A-dependent control of OGT. Under basal conditions, low glucocorticoid receptor (GR) activity maintains ALKBH5 expression at a modest level, resulting in high m^6^A methylation and degradation of *Ogt* mRNA. This limits OGT protein abundance and keeps gluconeogenic gene expression within a homeostatic range. In contrast, elevated glucocorticoid level enhances GR nuclear translocation and increases ALKBH5 expression, leading to reduced m^6^A on *Ogt* mRNA, OGT accumulation, and enhanced gluconeogenic gene expression, thereby driving hyperglycemia in T2DM. Pharmacological inhibition of ALKBH5 by 18l restores *Ogt* mRNA turnover and suppresses pathological gluconeogenesis. The figure was created using BioRender (https://biorender.com) under an Academic License

Previous studies have established that aberrant RNA m^6^A methylation is associated with diabetes [36, 37]. However, these studies primarily relied on bulk measurements of global m^6^A levels in peripheral blood from patients or whole liver tissues of diabetic models [18, 21, 38, 39]. While such approaches confirm overall m^6^A imbalance associated with T2DM, they provided limited insight into cell type-specific or mechanistic roles of m^6^A in disease progression. To address this, we employed transcriptome-wide m^6^A profiling in primary hepatocytes under pharmacological stimulation mimicking fasting-induced gluconeogenesis. This approach revealed a global reduction in m^6^A methylation, with hypomethylated RNAs enriched in pathways related to glycolipid metabolism, RNA processing, and autophagy, suggesting that m^6^A remodeling accompanies hepatocellular adaptation to increased gluconeogenic demand. Among the key m^6^A regulators examined, ALKBH5 was uniquely upregulated at both mRNA and protein levels in vitro and in vivo during gluconeogenic stimulation, suggesting its key role in mediating m^6^A remodeling in hepatocytes.

ALKBH5 has been implicated in diverse physiological and pathological processes, such as fertility [23, 40], neurodevelopment [41], metabolic homeostasis [42], and tumorigenesis [43–45]. Its expression and activity are subject to multilayered regulation, encompassing transcriptional regulation by factors such as HIF1α [46] and FOXD3 [47], as well as post-translational modifications like SUMOylation [48, 49], acetylation [50], and lactylation [51, 52]. Although ALKBH5 has been shown to respond to metabolic cues [53, 54], the upstream regulatory mechanisms controlling its expression remain elusive. Here, we demonstrate that glucocorticoid-induced GR activation transcriptionally upregulates ALKBH5 in hepatocytes, establishing a direct link between hormonal signals, the m^6^A machinery, and gluconeogenic gene expression. In contrast, although CREB has been reported as a transcriptional factor of ALKBH5 in ovarian cancer [55], pharmacological inhibition of CREB had no effect on ALKBH5 expression in hepatocytes (data not shown), underscoring tissue specificity of its transcriptional control. Moreover, activation of the glucagon-PKA signaling by Fsk did not alter ALKBH5 expression in hepatocytes, indicating that glucocorticoid and glucagon regulate ALKBH5 through distinct mechanisms. Together with recent findings that glucagon can modulate ALKBH5 activity via post-translational mechanism [54], these observations suggest that ALKBH5 is regulated at multiple levels during hepatic gluconeogenesis, enabling it to integrate diverse hormonal inputs.

Although aberrant m^6^A methylation has been implicated in the pathogenesis of T2DM [56, 57], the molecular substrates through which ALKBH5 influences hepatic glucose metabolism remain incompletely understood. Prior studies have largely focused on ALKBH5 in diabetes-related complications [53, 58, 59], which may not reflect its direct targets during gluconeogenic activation. A recent study reported that ALKBH5 regulates glucose homeostasis by demethylating *Gcgr* mRNA, thereby enhancing glucagon responsiveness [54]. In contrast, our work demonstrates that ALKBH5 regulates hepatic glucose production through a distinct substrate, *Ogt* mRNA, whose stabilization enhances OGT abundance. Given the well-established role of OGT-mediated O-GlcNAcylation in regulating gluconeogenic gene expression [33, 34], our findings position the ALKBH5–OGT axis as an upstream epitranscriptomic mechanism linking hormonal cues to hepatic gluconeogenesis. These findings highlight the substrate diversity of ALKBH5 and underscore the mechanistic complexity of m^6^A-mediated regulation in hepatic metabolism. Collectively, available evidence supports a model in which ALKBH5 coordinates hepatic glucose homeostasis through multiple, parallel downstream pathways.

The ubiquitous presence of m^6^A and its critical roles in diverse physiological and pathological processes position the m^6^A regulatory machinery as a compelling therapeutic target [60]. Consistent with this rationale, we demonstrate that 18l, a covalent small-molecule inhibitor of ALKBH5 [35], effectively reduces hepatic gluconeogenesis and improves glucose tolerance in diabetic mice, without apparent toxicity, providing a proof-of-concept for epitranscriptomic intervention in T2DM. In addition to ALKBH5, other components of the m^6^A network have emerged as potential targets for metabolic modulation [57]. For instance, METTL14 has been shown to post-transcriptionally regulate *G6pc*, thereby promoting gluconeogenesis in obese mice [61], whereas pharmacological inhibition of FTO with entacapone reduces body weight and fasting glucose through an FTO-FOXO1 axis in liver and adipose tissue [62]. Together, these findings delineate multiple nodes within the m^6^A regulatory network that converge on hepatic glucose production and systemic energy homeostasis, highlighting the broad translational potential of epitranscriptomic modulation as a therapeutic strategy for T2DM.

While our findings underscore the pivotal role of ALKBH5 in hepatic gluconeogenesis and the therapeutic potential of its inhibition, several limitations should be acknowledged. First, beyond *Gcgr* [54] and *Ogt*, our m^6^A-seq and RIP-seq analyses suggest the presence of additional metabolism-related targets, which requires further validation to define the broader downstream network of ALKBH5 in gluconeogenic control. Second, our study primarily relied on cell-based assays and mouse models, and future studies using human liver tissues will be required to assess the clinical relevance of ALKBH5-centered pathways. Finally, although 18l showed promising efficacy in vivo, its long-term safety profile, pharmacokinetics, and target specificity require thorough preclinical and ultimately clinical evaluation.

In summary, our work demonstrates dynamic m^6^A remodeling in hepatocytes during gluconeogenic stimulation and uncovers a GR-ALKBH5-OGT regulatory axis that links endocrine signaling to the m^6^A epitranscriptomic machinery. By integrating mechanistic dissection with pharmacological intervention, these findings expand our understanding of m^6^A regulation in hepatic glucose homeostasis and provide a conceptual framework for exploring epitranscriptomic strategies in the treatment of T2DM.

## Supporting information

List of Reagents and Resources.

Sequencing information of m6A-seq and RIP-seq.

GO enrichment analysis of genes with differential m6A methylation after Dex Fsk treatment.

GO enrichment analysis of genes with altered m6A methylation after Alkbh5 deletion.

GO enrichment analysis of genes corresponding to ALKBH5-bound RNAs identified by RIP-seq.

Hyper-methylated RNAs upon Alkbh5 deletion overlap with ALKBH5-binding RNAs.

## List of abbreviations

A3SS: alternative 3’ splice site
A5SS: alternative 5’ splice site
ActD: actinomycin D
Alb: albumin promoter
ALKBH5: AlkB homolog 5
AML12: Alpha Mouse Liver 12
ATAC: assay for transposase-accessible chromatin
CBC: complete blood count
cCREs: candidate cis-regulatory elements
CDS: coding sequence
cHET: conditional heterozygote
ChIP: chromatin immunoprecipitation
cKO: conditional knockout
CLUC: Cypridina luciferase
Con: control adenovirus
Cre: Cre-expressing adenovirus
Ctrl: control
Dex: dexamethasone
ENCODE: Encyclopedia of DNA Elements
ES: enrichment score
EV: empty vector
FC: fold change
FFPE: formalin-fixed paraffin-embedded
FOXD3: forkhead box D3
FPKM: fragments per kilobase of exon per million mapped reads
Fsk: forskolin
G6PC: glucose-6-phosphatase
GCGR: G-protein coupled receptor for glucagon
GLUC: Gaussia luciferase
GO: Gene Ontology
GR: glucocorticoid receptor
GTT: glucose tolerance test
H&E: hematoxylin and eosin
HCT: hematocrit
HCFC1: host cell factor C1
HECTD1: HECT domain E3 ubiquitin protein ligase 1
HGB: hemoglobin
HGP: hepatic glucose production
HIF1α: hypoxia-inducible factor 1-alpha
HMW: high molecular weight polysomes
hypo: hypomethylated
hyper: hypermethylated
IGV: Integrative Genomics Viewer
IHC: immunohistochemistry
LMW: low molecular weight polysomes
LYM: lymphocytes
m^6^A: N^6^-methyladenosine
LRP6: low density lipoprotein receptor-related protein 6
MCV: mean corpuscular volume
MPH: mouse primary hepatocytes
MXE: mutually exclusive exon
NEUT: neutrophils
O.D.: optical density
OE: overexpression
O-GlcNAcylation: protein O-linked β-N-acetylglucosamine modification
OGT: O-linked N-acetylglucosamine transferase
PCK1: phosphoenolpyruvate carboxykinase 1
PGC-1α: peroxisome proliferator-activated receptor gamma coactivator 1-alpha
PIKFYVE: phosphoinositide kinase, FYVE-type zinc finger containing
PLT: platelet count
PTT: pyruvate tolerance test
RBC: red blood cell count
RI: retained intron
RIP: RNA immunoprecipitation
RNF213: ring finger protein 213
RT-qPCR: reverse transcription-quantitative polymerase chain reaction
sc: scramble siRNA
SE: skipped exon
SUMOylation: small ubiquitin-like modifier conjugation
T2DM: type 2 diabetes mellitus
UCSC: University of California, Santa Cruz
UTR: untranslated region
Veh: vehicle
WBC: white blood cell count
WT: wild-type
YTHDF2: YT521-B homology domain family

## Acknowledgment

We thank Prof. Xuetao Cao and Yang Liu from Peking Union Medical College for kindly providing the pcDNA4-m*Alkbh5-*Myc-His plasmid. While ChatGPT (OpenAI) was utilized for minor grammatical fine-tuning and linguistic polishing to enhance clarity, the authors remain solely responsible for the study design, data analysis, and scientific interpretation.

## Author contributions

Y.C. performed study concept and design, formal analysis, investigation, and drafting, review, and revision of the manuscript. Y.F. contributed to data curation, formal analysis, and drafting, review, and revision of the manuscript. N.W. and X.X. performed investigation. Y.X., W.P., Q.W., Y.L., Q.L., A.L., and C.L. provided supporting investigation. H.W. and H.L. provided resources. C.L. performed study concept and design, funding acquisition, resource provision, and supervision. W.-M.T. performed study concept and design, funding acquisition, supervision, and manuscript review and revision. X.L. performed study concept and design, funding acquisition, supervision, and drafting, review, and revision of the manuscript. Y.N. performed study concept and design, funding acquisition, supervision, and drafting, review, and revision of the manuscript. All authors read and approved the final manuscript.

## Data Deposition

All data needed to evaluate the conclusions in this study are present in the paper and/or the Supplementary Materials. All the sequencing data presented in this study have been submitted to the Genome Sequence Archive database and are available under accession number CRA028572.

