## Supplementary Methods and Figures. for "ALKBH5 Promotes Hepatic Gluconeogenesis via m^6^A-mediated Stabilization of *Ogt* mRNA"

Yan Chen et al.

**This file includes:**

**Materials and methods**

**Supplementary Fig.1–6**

Supplementary Fig.1 m^6^A profiling of mouse primary hepatocytes treated with either DMSO or Dex/Fsk.

Supplementary Fig.2 Glucocorticoid receptor binds to *Alkbh5* DNA and enhances its transcription.

Supplementary Fig.3 Generation and phenotype analysis of hepatocyte-specific *Alkbh5* knockout (cKO) mice.

Supplementary Fig.4 m^6^A profiling analysis of *Alkbh5*-deficient mouse primary hepatocytes.

Supplementary Fig.5 m^6^A-mediated regulation of *Ogt* and OGT-dependent control of hepatic gluconeogenesis.

Supplementary Fig.6 The ALKBH5 inhibitor 18l modulates gluconeogenesis and demonstrates *in vivo* safety in mice.

**Supplementary Table 1–6 (See attached Excel files)**

Supplementary Table 1. List of Reagents and Resources.

Supplementary Table 2. Sequencing information of m^6^A-seq and RIP-seq.

Supplementary Table 3. GO enrichment analysis of genes with differential m^6^A methylation after Dex/Fsk treatment.

Supplementary Table 4. GO enrichment analysis of genes with differential m^6^A methylation after *Alkbh5* deletion.

Supplementary Table 5. GO enrichment analysis of genes corresponding to ALKBH5-bound RNAs identified by RIP-seq.

Supplementary Table 6. Hyper-methylated RNAs upon *Alkbh5* deletion overlap with ALKBH5- bound RNAs.

1. **Materials and methods**

**1.1. *In vitro* gene knockdown, knockout and overexpression**

For gene knockdown, MPH and AML12 cells were transfected with 40–80 nM siRNAs (GenePharma, Shanghai, China) using Rfect transfection reagent (Baidai biotechnology, 11013, Changzhou, China) according to the manufacturer’s protocol, and incubated for 48 h before subsequent analyses. Sequences of all siRNAs used in this study were included in Supplementary Table 1.

For gene knockout, MPH from *Alkbh5^flox/flox^* mice were allowed to adhere after plating and subsequently infected with Cre-expressing adenoviruses (WZ Biosciences, Shandong, China) at a multiplicity of infection (MOI) of 30. After 12 h of infection, the medium was replaced with fresh culture medium, and cells were maintained for another 36 h prior to collection for downstream analyses.

For gene overexpression, the mouse *Nr3c1*-overexpressing plasmid (pcDNA3.1-m*Nr3c1*-3×Flag-C) and its parental vector (pcDNA3.1-3×Flag-C) were purchased from Fenghui Biotechnology (Hunan, China). The mouse *Alkbh5*-overexpressing plasmid (pcDNA4-m*Alkbh5*-Myc-His) was kindly provided by Prof. Xuetao Cao and Yang Liu (IBMS/PUMC). The mouse *Ogt*-overexpressing plasmid (pcDNA3.1-m*Ogt*-Myc) was purchased from Tsingke Biotechnology (Beijing, China). The *Alkbh5* promoter reporter plasmids (pGL3.1-promoter-*Alkbh5* and pGL4.10-*Alkbh5*) and their corresponding empty vectors were obtained from General Biotechnology (Anhui, China). Plasmid transfections in MPH and AML12 cells were carried out using jetOPTIMUS® transfection reagent (Polyplus, 101000006, France) according to the manufacturer’s instructions. After 4 h of transfection, the medium was replaced with fresh culture medium, and cells were maintained for an additional 44 h prior to harvest for downstream analyses.

**1.2. RNA extraction and real-time quantitative PCR (RT-qPCR)**

Mouse liver tissues or cultured cells were homogenized in TRIzol reagent, and total RNA was extracted following the manufacturer’s instructions. Reverse transcription was performed using ReverTra Ace qPCR RT Master Mix (TOYOBO, FSQ-301, Japan) according to the manufacturer’s protocol. Quantitative PCR was carried out in triplicate using SYBR® qPCR Mix (TOYOBO, QPS-201, Japan) on a real-time PCR system. Relative gene expression was calculated using *18S* rRNA or *U1* snRNA as the internal control. Primer sequences are listed in Supplementary Table 1.

**1.3. N^6^-methyladenosine (m^6^A) sequencing (m^6^A-seq), m^6^A immunoprecipitation with quantitative PCR (m^6^A-IP-qPCR), and data analysis**

Total RNA was extracted from primary mouse hepatocytes using TRIzol reagent. For each reaction, 12 µg RNA was supplemented with 3.6 µL of synthetic spike-in RNAs consisting of equal amounts of m^6^A-modified Gaussia luciferase (*Gluc*) and unmodified Cypridina luciferase (*Cluc*) RNAs (New England Biolabs, E1610S, USA) and the RNA was fragmented at 95 °C for 40 s to yield 100–150 nt fragments using RNA Fragmentation Reagents (Thermo Fisher Scientific, AM8740, USA). For each immunoprecipitation (IP) reaction, 10 µg of fragmented RNA was incubated with 1.5 µg of anti-N^6^-methyladenosine antibody (Synaptic Systems, 202003, Germany) overnight at 4 °C, followed by incubation with 20 µL of Dynabeads Protein A (Thermo Fisher Scientific, 10009D, USA) for 2 h at 4 °C. Beads were washed twice with m^6^A -containing buffer (Berry & Associates, PR3732, USA) for 1 h each, and RNA was eluted and purified using the RNeasy MinElute Cleanup Kit (Qiagen, 74204, Germany). For library preparation, 10 ng of input RNA and 20 ng of m^6^A-IP RNA were used with the SMARTer Stranded Total RNA-seq Kit v2 (Takara, 634413, Japan). Library concentrations were measured with the Qubit dsDNA HS Assay Kit (Invitrogen, Q32851, USA), and sequencing was performed on an Illumina NovaSeq 6000 platform with paired-end 150 bp reads.

For data processing, we used FastQC (v0.11.3) [1] to assess the quality of raw sequencing data. Trimmomatic (v0.36) [2] was employed for adapter trimming, removal of low-quality bases (bases with quality scores < 3), and exclusion of reads shorter than 50 bp. Cleaned high-quality paired-end reads were then aligned to the mouse reference genome (GRCm38) using HISAT2 (v2.1.0) [3]. Only uniquely mapped reads with a mapping quality score greater than 20 were retained for downstream analysis. Aligned reads were sorted and filtered using SAMtools (v1.9) [4], and exon-level read counts were quantified using the featureCounts from the Subread R package (v1.32.4) [5] with gene annotations provided by Ensembl GRCm38.86. To visualize genome-wide read distribution, we generated alignment tracks with IGVtools (v2.4.17) [6]. Final images were annotated using Adobe Illustrator 2024.

The identification of m^6^A-modified peaks was performed using the exomePeak R package (v2.16.0) [7], with the corresponding Input sample as the control. To quantify read enrichment at m^6^A peak summits, we processed IP and Input BAM files using BEDTools (v2.26.0) [8]. Peaks identified by exomePeak were first sorted, and base-level coverage was computed using bedtools coverage -d. For each peak, the position with the highest read depth was defined as the summit, and a ±75 bp window (151 bp total) centered at the summit was generated. To ensure peak relevance, summit regions were intersected with BAM files using a strand-specific approach and a minimum 50% overlap threshold (-F 0.50). Final read counts over these regions were calculated for both IP and Input samples using multiBamCov. These values were subsequently used for enrichment quantification and visualization.

For enrichment analysis, FPKM (fragments per kilobase of exon per million mapped reads) values were calculated for both IP and Input samples. The Enrichment Score (ES) for each summit region was then calculated using the following formulas:

$$\mathrm{FPKM}_{\mathrm{IP}}=\frac{\mathrm{RC}_{\mathrm{IP}}\times{10}^{9}}{\mathrm{TRC}_{\mathrm{IP}}\times peak length}$$

$$\mathrm{FPKM}_{\mathrm{Input}}=\frac{\mathrm{RC}_{\mathrm{Input}}\times{10}^{9}}{\mathrm{TRC}_{\mathrm{Input}}\times peak length}$$

$$ES=\frac{\mathrm{FPKM}_{\mathrm{IP}}}{\mathrm{FPKM}_{\mathrm{Input}}}$$

where RC_IP_ and RC_Input_ represent the read counts mapped to the peak regions in the IP and Input samples, respectively, and TRC_IP_ and TRC_Input_ refer to the total read counts mapped to the exonic regions in each sample.

Peaks were considered high confidence if they met all the following criteria: 1) false discovery rate (FDR) < 0.05; 2) FPKM_IP_ > 2; 3) and ES > 1.5. Peak annotation was performed using Ensembl gene annotation, and custom Perl scripts were employed to assign peaks to genomic features. Motif discovery was conducted using HOMER (v4.10.3) [9] using ±75 bp sequences centered on m^6^A peak summits. Peak regions were derived by intersecting read depth data with high-confidence peaks using BEDTools and processed with custom scripts to retain strand information. The mouse reference genome was used, with motif lengths set to 5 and 6 nucleotides. The Guitar R package (v2.20.0) [10] was used to assess the positional distribution of peaks along mRNA transcripts relative to gene structures. To visualize enrichment patterns and compare signal intensities across conditions, boxplots and cumulative distribution plots were generated using ggplot2 R package (v3.5.2) [11]. All statistical tests, unless otherwise specified, were performed using a two-tailed Student’s *t*-test.

Differential m^6^A methylation analysis was performed using the exomePeak R package. To identify differentially methylated m^6^A-modified regions between treated and control conditions, overlapping peaks were defined as shared m^6^A regions if they exhibited at least 50% overlap of either peak in the treated or control group. For each shared peak, FPKM values were calculated for both IP and Input samples, along with the ES. Differentially methylated peaks were classified based on the specific criteria: hypermethylated peaks were defined by FDR < 0.05, log_2_(FoldChange) > 0.58, and ES in the treated group > 1.5, while hypomethylated peaks were characterized by FDR < 0.05, log_2_(FoldChange) < -0.58, and ES in the control group > 1.5. Volcano plots were generated using ggplot2 to visualize differential m^6^A methylation patterns.

For m^6^A-IP-qPCR, total RNA was isolated from MPH. For each reaction, 12 µg of fragmented RNA was supplemented with 3.6 µL of synthetic spike-in RNAs consisting of equal amounts of m^6^A-modified *Gluc* and unmodified *Cluc* RNAs and fragmented to 300–400 nt at 85 °C for 30 s. 10 µg of the fragmented RNA was subjected to immunoprecipitation using 2.5 µL (2.3 µg) of anti-m^6^A antibody (Abcam, ab151230, UK) and 20 µL of Dynabeads Protein A. The immunoprecipitation and washing steps were carried out as described for the m^6^A-seq procedures. Recovered RNA, together with corresponding input samples, was subjected for RT-qPCR. The primer sequences used for gene-specific detection are listed in the Supplementary Table 1.

**1.4. Protein extraction and Western blotting analysis**

Mouse liver tissues or cultured cells were washed with cold PBS and lysed in RIPA buffer supplemented with protease and phosphatase inhibitor cocktails (Roche, 04693159001 and 04906837001, Switzerland). Lysates were clarified by centrifugation, and protein concentrations were determined using a BCA Protein Assay Kit (GenStar, E162-05, Beijing, China) and quantified by detecting the absorbance at 562 nm. Equal amounts of protein (20–50 μg) were separated on SDS-PAGE gels and transferred onto 0.45 μm PVDF membranes (Millipore, IPVH00010, Germany). Membranes were blocked in 5% non-fat milk in Tris-Buffered Saline with 0.05% Tween-20 for 1 h at room temperature and incubated overnight at 4 °C with primary antibodies. After washing with TBST, membranes were incubated with HRP-conjugated secondary antibodies for 1 h at room temperature. Protein signals were detected using Clarity ECL substrate (Bio-Rad, 170-5060, USA) and visualized with a gel imaging system (Tanon, 5800, Shanghai, China). Quantification of band intensities was performed using ImageJ (v1.54g). Detailed information of the antibodies used is listed in Supplementary Table 1.

**1.5. Immunohistochemistry (IHC) analysis**

Immunohistochemical staining was performed on formalin-fixed paraffin-embedded (FFPE) liver tissue sections. Briefly, 4 μm-sections were deparaffinized in xylene and rehydrated through graded ethanol. Antigen retrieval was carried out by microwave heating in citrate buffer (pH 6.0) for 15 min, followed by cooling to room temperature. Endogenous peroxidase activity was quenched by incubation with 3% hydrogen peroxide (ZSGB-Bio, ZLI-9311, Beijing, China) in PBS for 10 min. The sections were then blocked with 2.5% goat serum for 30 min at room temperature and incubated overnight at 4 °C with primary antibody against ALKBH5 (Sigma-Aldrich, HPA007096, 1:400, USA). After rinsing with PBS, slides were incubated with HRP-conjugated secondary antibody for 45 min at room temperature, visualized using DAB substrate (Vector, SK-4100, USA) for 1.5 min, and counterstained with hematoxylin. Finally, the slides were scanned with a Panoramic MIDI II digital slide scanner (3D Histech, Hungary) for image acquisition and analysis.

**1.6. Chromatin Immunoprecipitation (ChIP) assay**

Chromatin immunoprecipitation was performed using the SimpleChIP® Plus Enzymatic Chromatin IP Kit (Magnetic Beads) (Cell Signaling Technology, 9005, USA) following the manufacturer’s protocol with minor modifications. AML12 cells were seeded into 15-cm culture dishes and grown to approximately 80–90% confluency. Cells were transfected with the pcDNA3.1-m*Nr3c1*-3×Flag-C. At 24 h post-transfection, the culture medium was replaced with DMEM/F12 supplemented with 2% FBS containing 1 μM dexamethasone and 10 μM forskolin for 24 h. Following drug treatment, cells were cross-linked with 1% formaldehyde for 10 min at room temperature, and the reaction was quenched by adding 2 mL of 10× glycine per dish for 5 min at room temperature. Cells were collected, nuclei were isolated, and chromatin was digested with micrococcal nuclease to obtain fragments of 150–900 bp.

For each IP, 500 μL of diluted chromatin was incubated with antibody at 4 °C for 4 h or overnight, followed by the addition of 30 μL of ChIP-Grade Protein G magnetic beads (Cell Signaling Technology, 9005, USA) for 2 h at 4 °C. After washing, protein–DNA complexes were eluted, cross-links were reversed with Proteinase K (Cell Signaling Technology,10012, USA) at 65 °C, and DNA was purified with spin columns (Cell Signaling Technology,10010, USA). The recovered DNA was analyzed by qPCR using primers for *Rpl30* (positive control, provided with the kit) and for *Alkbh5* (listed in Supplementary Table 1).

**1.7. Dual-luciferase reporter assay**

Cells were plated in 6-well plates and co-transfected with three plasmids: pcDNA3.1-m*Nr3c1* plasmid (2 µg), pGL4.10-*Alkbh5* or pGL3-promoter-*Alkbh5* reporter plasmid (0.5 µg), and pRL-TK control plasmid (0.05 µg). 24 h later, cells were treated with culture medium with 2% FBS containing 5 µM Dex for an additional 24 h. After treatment, cells were washed twice with PBS and lysed with 250 µL of cell lysis buffer from the Dual-Luciferase Reporter Assay Kit (Beyotime, RG027, Shanghai, China) on a shaker for 20 min. Cell lysates were collected by pipetting, and debris was removed by centrifugation at 13,300 rpm for 5 min at 4 °C. Luciferase activities were measured by first adding the Luciferase Assay Reagent to detect firefly luciferase, followed by Stop & Glo reagent to quench firefly activity and initiate Renilla luciferase measurement. Firefly luciferase activity was normalized to Renilla luciferase activity to account for variations in transfection efficiency and cell viability.

1.8. **Glucose output assay**

MPH were seeded into 6-well plates and transfected with siRNAs or plasmids as previously described. 30 h later, the cells were washed and cultured in glucose- and serum-free RPMI-1640 medium (Pricella, PM150130, Beijing, China) for 5 h. Following the starvation period, the medium was replaced with glucose-free RPMI-1640 medium containing 1 μM Dex, 10 μM Fsk, 20 mM sodium L-lactate (Aladdin, S108838, Shanghai, China), and 2 mM sodium pyruvate (Solarbio, P8380, Beijing, China). After 12 h, supernatant was collected and measured for glucose concentration by using a Glucose Content Assay Kit for Liquid Samples (Pulilai Biotech, E1010, Beijing, China).

For normalization, the total cellular protein concentration was determined. The cells were digested with 0.25% trypsin (Thermo Fisher Scientific, C25200056, USA), and the cell pellets were collected, washed with PBS, and lysed with 200 μL of RIPA lysis buffer. The protein concentration was then measured using a BCA Protein Assay Kit. The glucose content was normalized to the total cellular protein concentration for each well.

**1.9. Pyruvate tolerance test (PTT) and glucose tolerance test (GTT)**

For pyruvate tolerance tests (PTT), male *db/db* mice (7–8 weeks old) were fasted for 16 h and injected intraperitoneally with sodium pyruvate at 2.5 g/kg (injection volume: 6 μL/g body weight). For glucose tolerance tests (GTT), *db/db* mice were fasted for 16 h prior to intraperitoneal injection of glucose at 2 g/kg (injection volume: 3 μL/g body weight). For non-diabetic mice, including 5­6 months male *Alkbh5*-KO and cKO mice, their littermate controls, and 2 months male WT mice treated with ALKBH5 inhibitor 18l, both PTT and GTT were performed following a 6 h fasting period. For PTT, mice received sodium pyruvate at 2.5 g/kg (injection volume: 6 μL/g body weight). For GTT, mice received intraperitoneal glucose at 2 g/kg (injection volume: 6 μL/g body weight). Blood glucose concentrations were measured from tail vein blood at 0, 15, 30, 60, 90, and 120 min post-injection using a OneTouch Verio Vue® glucometer (USA). Sample size determination for each experimental group, as indicated in the figure legends (ranging from n=4 to n=8), was determined based on our pilot study data and established practices in the field for *in vivo* metabolic studies. This sample size was selected to ensure adequate statistical power to detect biologically relevant differences while adhering to the principles of reducing the number of animals used. Specifically, we used a minimum of n=6 for key in vivo studies (e.g., PTT/GTT, drug efficacy) to account for inherent biological variability in mouse models such as *db/db* mice, thereby minimizing the risk of Type II errors.

**1.10. RNA-immunoprecipitation (RIP) -seq, data analysis and RIP-qPCR**

MPH (~5 × 10^7^ cells) were lysed in nondenaturing lysis buffer (50 mM Tris-HCl, pH 7.4, 250 mM NaCl, 0.5% Triton X-100, 1 mM DTT, 2 mM EDTA, 1 mM NaF, 1× protease inhibitor cocktail, and 0.04 U/μL RNasin (Promega, N2515, USA) at 4 °C. For each RIP reaction, 10 mg of protein was incubated with antibody at a final concentration of 5 μg/mL for 6 h at 4 °C, with the remaining lysate reserved as input. Antibody–protein complexes were subsequently captured by incubation with 45 μL of Dynabeads Protein A, pre-blocked with BSA protein standard (2 mg/mL), overnight at 4 °C with gentle rotation.

On the next day, 1/10 of the beads were subjected to Western blotting analysis to verify immunoprecipitation efficiency. The remaining complexes were digested with proteinase K (Sigma-Aldrich, P2308, Germany) at 55 °C, 1,100 rpm for 30 min, and the RNA was purified from both input and IP fractions by TRIzol reagent.

For RIP-seq, RNA was extracted from input (10 ng), IP (20 ng), and IgG control (25 ng) samples and subjected to library preparation following the same protocol as described for m^6^A-seq, followed by sequencing on Illumina NovaSeq 6000. Data processing followed the procedures described for m^6^A-seq.

To identify transcripts bound by ALKBH5, we performed peak calling using MACS2 (v2.2.7.1) [12]. Specifically, IP vs Input and IgG vs Input comparisons were conducted separately with the callpeak function. To identify differentially enriched peaks between the IP and IgG samples, we utilized the bdgdiff module in MACS2, which computes differential enrichment based on signal intensity across conditions. The resulting peaks were filtered and annotated using the corresponding aligned BAM files generated by HISAT2, from which read counts were extracted for each peak using bedtools multicov. FPKM values were then calculated for each peak to quantify the binding strength of ALKBH5. FPKM normalization accounts for both the length of each peak and the total number of mapped reads. Then, we obtain transcripts that ALKBH5 bound by annotating peaks with the reference genome. Transcripts were considered significant based on the following criteria: score > 5, FPKM_IP_ ≥ 15, log_2_(FoldChange _IP vs Input_) > 3, log_2_(FoldChange _IP vs IgG_) > 2.

For RIP-qPCR, a total of 100 ng RNA was used for reverse transcription, and qPCR was performed with primers listed in Supplementary Table 1.

**1.11. Subcellular fractionation analysis of RNA**

MPH were collected and subjected to subcellular fractionation using the PARIS™ Kit (Thermo Fisher Scientific, AM1921, USA) according to the manufacturer’s instructions. Briefly, cells were lysed in Cell Disruption Buffer, and the cytoplasmic and nuclear fractions were separated by centrifugation. Total RNA from each fraction was purified and dissolved in equal volumes of nuclease-free water. For downstream analysis, equal volumes of nuclear and cytoplasmic RNA solutions were subjected to RT-qPCR. *Gapdh* was used as a cytoplasmic marker and *U1* as a nuclear marker. Primer sequences used in this study are listed in the Supplementary Table 1.

**1.12. RNA stability assay**

MPH were seeded into 6-well plates and subjected to either plasmid or siRNA transfection for 40 h. Transcription was subsequently inhibited by adding 1 mL of complete RPMI 1640 medium containing 5 μg/mL Actinomycin D (Sigma-Aldrich, A9415, Germany) to each well. Cells were harvested at 0, 3, 6, and 9 h after Actinomycin D treatment, or at other indicated time intervals. Total RNA was isolated using the TRIzol reagent, followed by RT-qPCR. *18S* rRNA or *U1* snRNA was used as an internal control. RNA decay rate was analyzed according to the following exponential decay model:

N_t_/N_0_=e^−kt^

where t is the duration of transcriptional inhibition, N_0_ is the RNA abundance at time 0, and N_t_ is the abundance at time t. The half-life (T_1/2_) of RNA was calculated as：

T_1/2_=ln (2)/k

Primer sequences used for qPCR are listed in the Supplementary Table 1.

**1.13. Polysome profiling analysis**

AML12 cells were seeded into 10-cm dishes and transfected with pcDNA4-m*Alkbh5*-Myc-His for 24 h, then re-plated into 15-cm dishes and harvested 48 h later. Approximately 2 × 10^7^ cells were collected per sample. Before harvest, cells were washed with ice-cold PBS, briefly trypsinized, and treated with culture medium containing 100 μg/mL cycloheximide (CHX, MCE, HY-12320, Shanghai, China) at 37 °C for 8 min with gentle mixing. After centrifugation, the pellet was lysed on ice in polysome lysis buffer (20 mM Tris-HCl pH 7.4, 150 mM NaCl, 5 mM MgCl_2_, 1% Triton X-100, 100 μg/mL CHX, 1 mM DTT, RNase inhibitor, and protease inhibitor cocktail) for 30 min, followed by clarification at 12,000 × g for 10 min at 4 °C. A portion of the supernatant was reserved as input. The remaining lysates were loaded onto 20–50% sucrose gradients prepared in the same buffer and centrifuged at 4 °C for 3 h. Gradients were fractionated using a Gradient Fractionator (BioComp, Canada) into 37 fractions. For quality control, 40 μL of fractions 8–14 and 21–26 together with the remaining material were analyzed by Western blot. RPS6 was used as the marker for 40S/80S, RPL11 for 60S/80S, and RPL26 for polysomes. For RNA analysis, RNA from individual fraction from 8–30 was extracted with TRIzol, dissolved in 20 μL of water, and reverse transcribed using equal RNA volumes (2 μL). The abundance of *Ogt* mRNA in each fraction was measured by qPCR and normalized to input.

**1.14. Cell viability assay**

ALKBH5 inhibitor 18l was kindly provided by Prof. Hong Liu (SIMM/CAS). MPH and AML12 cells were seeded into 96-well plates at an appropriate density to reach approximately 70–80% confluency at the time of treatment. After adherence, the culture medium was replaced with RPMI-1640 or DMEM/F12 supplemented with 2% FBS to reduce cell proliferation. 18l was dissolved in DMSO and applied at concentrations of 0, 2.5, 5, 10, and 15 µM. The DMSO content was equalized across all wells by supplementing with vehicle to maintain consistent solvent conditions. Cells were incubated with the drugs for 48 h. Following treatment, cell viability was assessed using the CCK-8 assay kit (Beyotime, C0042, Shanghai, China) according to the manufacturer’s instructions. Briefly, 10 µL of CCK-8 solution was added to each well containing 100 µL of culture medium, and cells were incubated at 37 °C for 1–2 h. Absorbance at 450 nm was measured using a microplate reader. Relative cell viability was calculated by normalizing the absorbance values to vehicle-treated control cells.

**1.15. *In vitro* and *in vivo* application of 18l**

*In vitro*, MPH and AML12 cells were seeded into 6-well culture plates at an appropriate density to achieve approximately 70–80% confluency at the time of treatment. 18l were dissolved in DMSO to prepare stock solutions and diluted in culture medium to final working concentrations of 5–10 µM. Cells were exposed to the treatment for 48 h.

For *in vivo* application, 18l was dissolved in a solvent mixture consisting of 5% DMSO, 10% ethanol (Sinopharm, 10009218, Beijing, China), 40% PEG400 (Solarbio, IP9000, Beijing, China), and 45% saline. The solution was sterilized through a 0.22-μm filter, aliquoted, and stored at −80 °C until use. For pharmacological evaluation, 8-week-old male C57BL/6J mice were randomly assigned to two groups using the RAND function in Microsoft Excel: a control group receiving vehicle and an experimental group receiving 18l at 5 mg/kg dose. 18l was administered via intraperitoneal injection once daily for 14 consecutive days. Body weights and tolerance tests were monitored throughout the treatment period. At the end of the period, mice were euthanized for safety assessment and subsequent analyses. Whole blood was collected for a complete blood count (CBC), which was performed at the Department of Pathology, Peking Union Medical College Hospital. For serum analysis, blood samples were collected and allowed to clot at room temperature for 30 min before being centrifuged at 3,000 rpm for 15 min to obtain serum. Serum levels of alanine aminotransferase (ALT) and aspartate aminotransferase (AST) were measured as indicators of liver function, also at the Department of Pathology, Peking Union Medical College Hospital. For histological analysis, major organs, including the heart, liver, spleen, lung, and kidney, were surgically excised. A portion of each organ was immediately fixed in 4% paraformaldehyde (PFA) for 24 h. The fixed tissues were then processed, embedded in paraffin, and sectioned into 4-μm-thick slices. These sections were subsequently stained with hematoxylin and eosin (H&E) for morphological examination. 7–8-week-old male *db/db* mice were randomly assigned to a control group or an experimental group receiving 18l at 2 mg/kg via intraperitoneal injection once daily. Body weights and tolerance tests were monitored daily, and mice were sacrificed at the end of the treatment period for tissue collection and subsequent analyses. Due to the necessity of accurate drug administration, the personnel responsible for preparing and injecting the treatment and vehicle solutions were aware of the group allocation. To control for bias, the objectivity of the blood glucose concentration readings (PTT and GTT) was relied upon, while all subsequent subjective assessments, including tissue processing, H&E staining assessment, and statistical analyses, were strictly performed by investigators blinded to the group identity.

**1.16. Alternative splicing analysis**

Alternative splicing events were analyzed using rMATS (v4.1.2) [13], based on m^6^A-seq Input samples from *Alkbh5* knockout experiments. To visualize specific splicing events, the rMATS2 sashimiplot (v2.0.2) was used to generate sashimi plot*.* A pie chart depicting the proportions of different types of splicing events was generated using the ggplot2 R package.

**1.17. Gene Ontology enrichment analysis**

For each cell type, Gene Ontology (GO) enrichment analysis was performed to assess biological processes, molecular functions, and cellular components associated with corresponding gene sets, using the clusterProfiler R package (v4.6.2) [14]. GO terms were considered statistically significant if both the *p*-value and *q*-value were less than 0.05.

**1.18. Differential gene expression analysis**

DESeq2 (v1.44.0) [15] was used to perform differential gene expression analysis to compare the expression of known m^6^A regulators in response to Dex/Fsk treatment. Two public datasets GSE130525 [16] and GSE72086 [17] from GEO database were used to compare expressions changes of known m^6^A regulators under fasting conditions. For GSE130525, DESeq2 was used to perform differential gene expression analysis. Gene expression was normalized as log_2_(FPKM+1) and standardized by *z*-score. Since GSE72086 didn’t include raw counts data, we performed differential expression analysis using log_2_(FPKM+1) value. Log_2_(FoldChange) was calculated as the mean difference between groups, and *p*-value were adjusted using the Benjamini-Hochberg method. Expression values were *z*-score normalized across samples for heatmap visualization. ComplexHeatmap R package (v2.20.0) [18] was used to generate heatmaps.

**1.19. ChIP-seq data analysis**

To explore the potential upstream regulatory mechanism of *Alkbh5*, we utilized UCSC Genome Browser (mouse mm10 assembly) [19]. Visualization tracks included ReMap ChIP-seq data [20] and open chromatin ATAC-seq signals from day 0 (P0) mouse liver, obtained from ENCODE3 (UCSD/Ren) [21-23]. Genomic coordinates were annotated using the NCBI Refseq database (release 2020-07-09) [24]. Information on candidate cis-regulatory elements (cCREs) was obtained from the ENCODE project [25], with data last updated on the UCSC genome browser on 2021-05-26. We examined ChIP-seq data from public dataset GSE137978 [26] in UCSC Genome Browser, and we visualize the genome-wide read distribution of another ChIP-seq analysis data from GSE72087 [17] by IGVtools.

**1.20. Statistical analysis**

The data are presented as the means ± SEMs. The number of experimental repeats is detailed in the figure legends. Statistical analyses were conducted via GraphPad Prism (v10.4.1). Comparisons were performed via two-tailed unpaired Student’s t-test, one-way ANOVA or two-way ANOVA, as specified in the figure legends. A *p*-value of less than 0.05 was considered statistically significant. Data points for Pyruvate Tolerance Test (PTT) and Glucose Tolerance Test (GTT) were excluded only if there was clear evidence of procedural failure (e.g., instrument malfunction, observable dosing error) or if they met objective statistical criteria for outliers (e.g., more than 2 standard deviations from the mean) as pre-specified in the statistical analysis plan. The reasons for any data exclusion are reported.

**Supplementary figures**

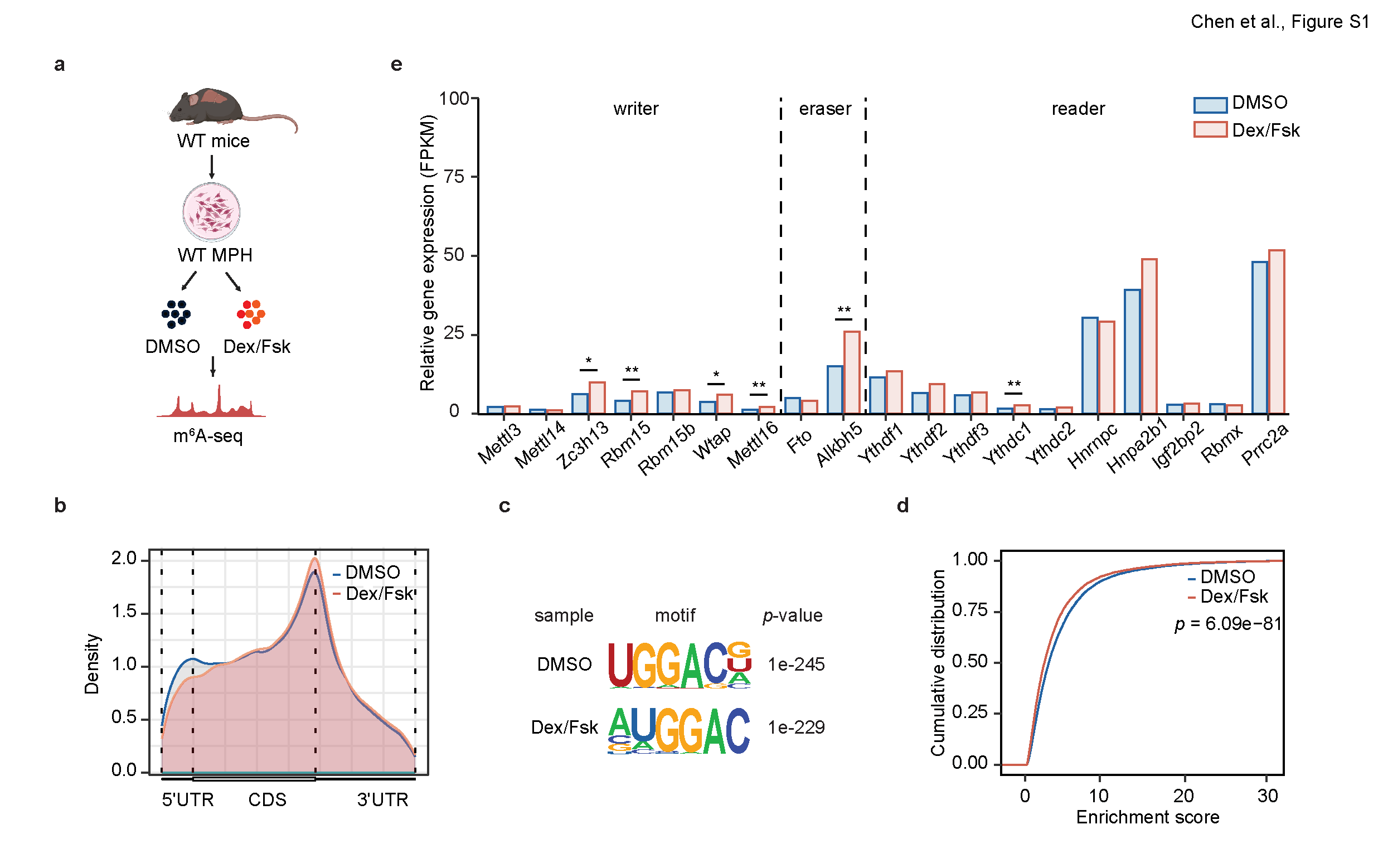

**Fig. S1 m^6^A profiling of mouse primary hepatocytes treated with either DMSO or Dex/Fsk. Related to Fig. 1.**

**a** Schematic workflow of m^6^A-seq performed on MPH treated with either DMSO or Dex/Fsk. **b** Metagene profiles showing the distribution of m^6^A peaks across mRNA transcripts. **c** Top consensus motif identified from m^6^A peaks. **d** Cumulative distribution of enrichment scores showing the relative methylation levels of all m^6^A peaks. **e** Relative RNA expression levels of m^6^A regulators. Data are represented as mean ± SEM. Statistical significance was calculated by two-tailed unpaired Student’s t-test (**d**) or Wald test (DESeq2, E). **p* < 0.05, ***p* < 0.01, and ****p* < 0.001.

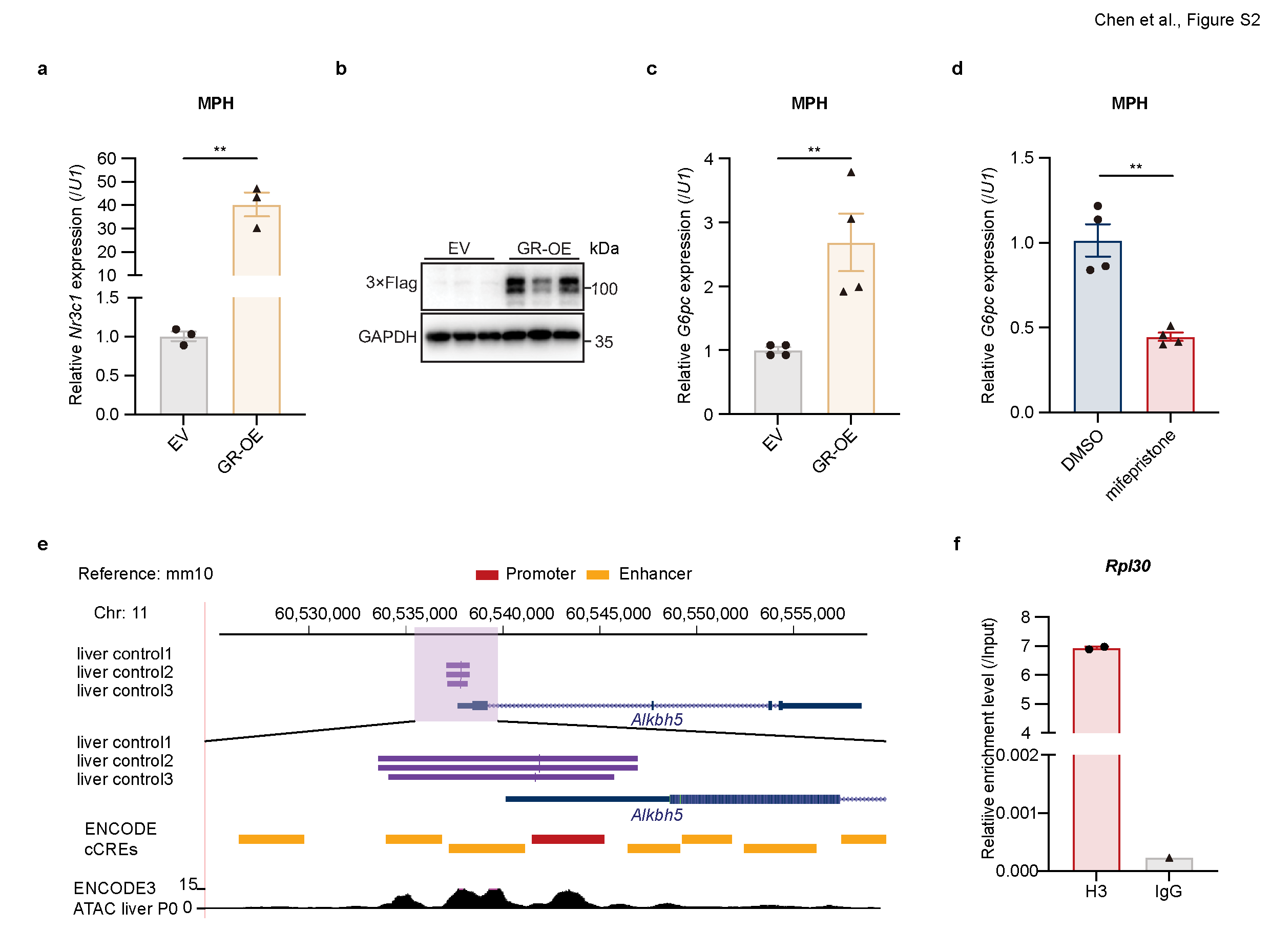

**Fig. S2 Glucocorticoid receptor binds to *Alkbh5* DNA and enhances its transcription. Related to Fig. 2.**

**a** RT-qPCR analysis of *Nr3c1* expression in GR-overexpressing (OE) MPH (n = 3). **b** Representative Western blot of GR expression in GR-OE MPH (n = 3). **c** RT-qPCR analysis of *G6pc* expression in GR-OE MPH as a positive control (n = 4). **d** RT-qPCR analysis of *G6pc* expression in GR inhibitor mifepristone-treated MPH (n = 4). **e** UCSC Genome Browser snapshot depicting the GR ChIP-seq profile at the *Alkbh5* gene locus, alongside candidate cis-regulatory elements (cCREs) annotated by the ENCODE project, generated from the publicly available dataset GSE137978. **f** ChIP-qPCR results in AML12 cells showing enrichment of Histone H3 at the *Rpl30* gene, with IgG serving as a negative control. Enrichment was calculated relative to input. Data are represented as mean ± SEM. Statistical significance was calculated by two-tailed unpaired Student’s t-test (**a**, **c**, **d**). **p* < 0.05, ***p* < 0.01, and ****p* < 0.001.

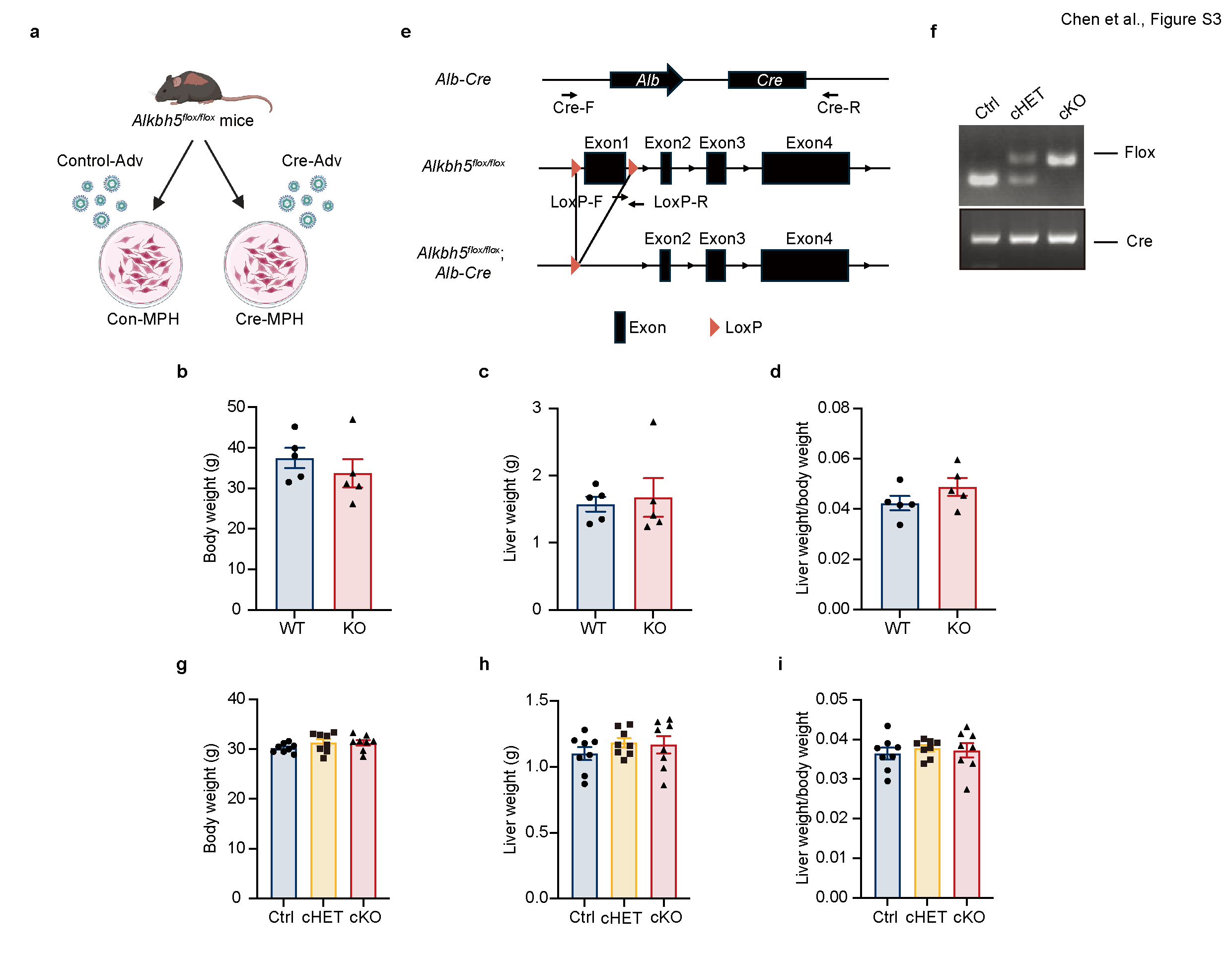

**Fig. S3 Generation and phenotype analysis of hepatocyte-specific *Alkbh5* knockout (cKO) mice. Related to Fig. 3.**

**a** Schematic of the workflow for generating *Alkbh5*-knockout MPH. **b–d** Body weight (**b**), liver weight (**c**), and liver weight-to-body weight ratio (**d**) in WT and *Alkbh5*-KO mice (n = 5). **e** Schematic of the workflow for generating hepatocyte-specific *Alkbh5*-knockout mice by crossing *Alkbh5^flox/flox^* mice with *Alb*-*Cre* transgenic mice. **f** Representative PCR genotyping results of hepatocyte-specific *Alkbh5* knockout mice. **g–i** Body weight (**g**), liver weight (**h**), and liver weight-to-body weight ratio (**i**) in Ctrl, hepatocyte-specific *Alkbh5*-cHET and cKO mice (n = 8). Data are represented as mean ± SEM. Statistical significance was calculated by two-tailed unpaired Student’s t-test (**b**–**d**) or one-way ANOVA with Dunnett’s multiple comparisons test (**g**–**i**). **p* < 0.05, ***p* < 0.01, and ****p* < 0.001.

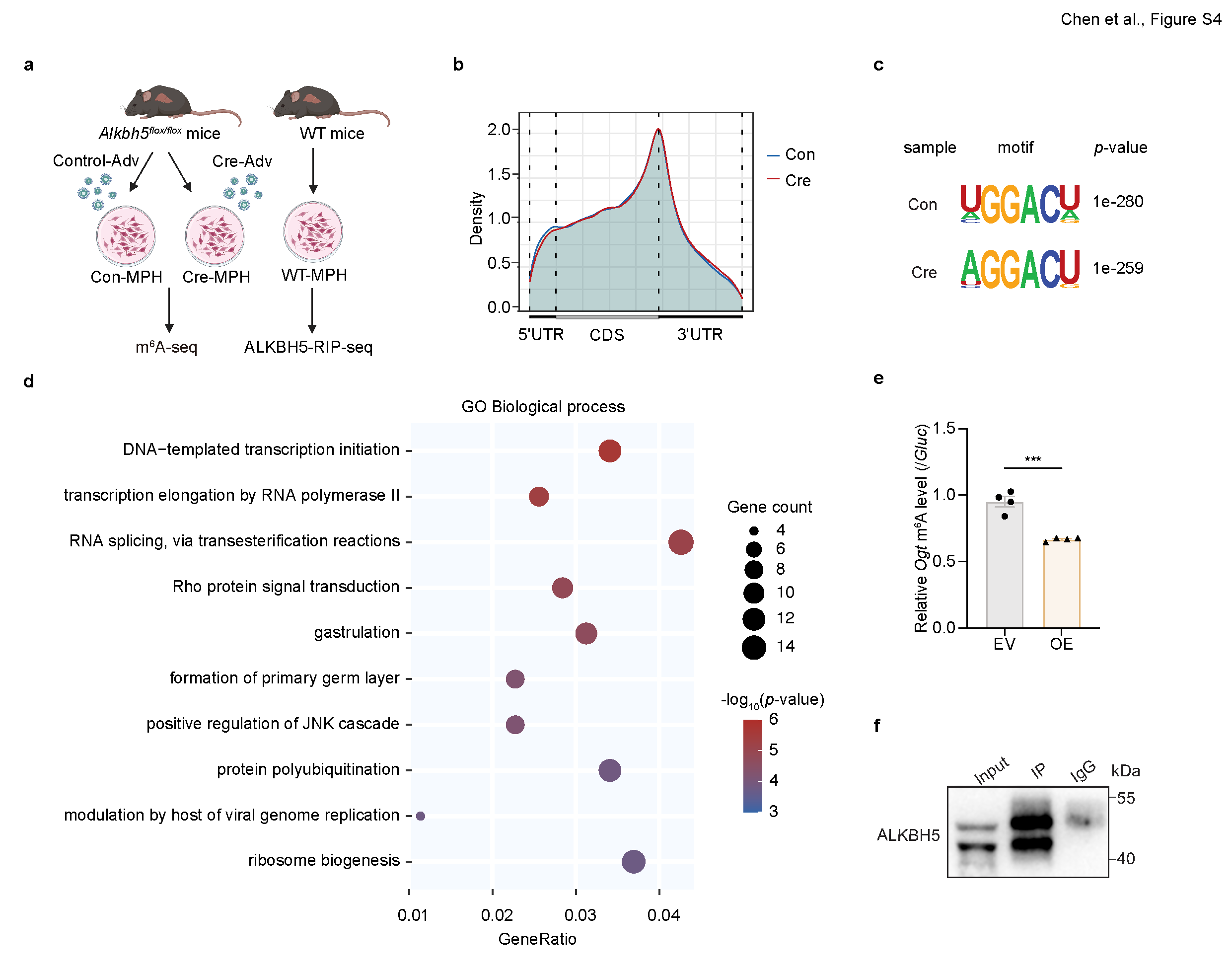

**Fig. S4 m^6^A profiling analysis of *Alkbh5*-deficient mouse primary hepatocytes. Related to Fig. 4.**

**a** Schematic workflow of m^6^A-seq and ALKBH5-RIP-seq performed in MPH. **b** Metagene profiles showing the distribution of m^6^A peaks across mRNA transcripts in MPH infected with control (Con) or Cre-expressing (Cre) adenovirus. **c** Top consensus motif identified from the m^6^A peaks. **d** GO analysis of genes with decreased RNA m^6^A methylation in *Alkbh5*-deficient MPH. **e** m^6^A-IP-qPCR results showing the effect of ALKBH5 overexpression on the m^6^A levels of *Ogt* transcript in MPH (n = 4). **f** Representative Western blot showing the efficiency of ALKBH5 immunoprecipitation (IP) in MPH. Data are represented as mean ± SEM. Statistical significance was calculated by two-tailed unpaired Student’s t-test (**e**). **p* < 0.05, ***p* < 0.01, and ****p* < 0.001.

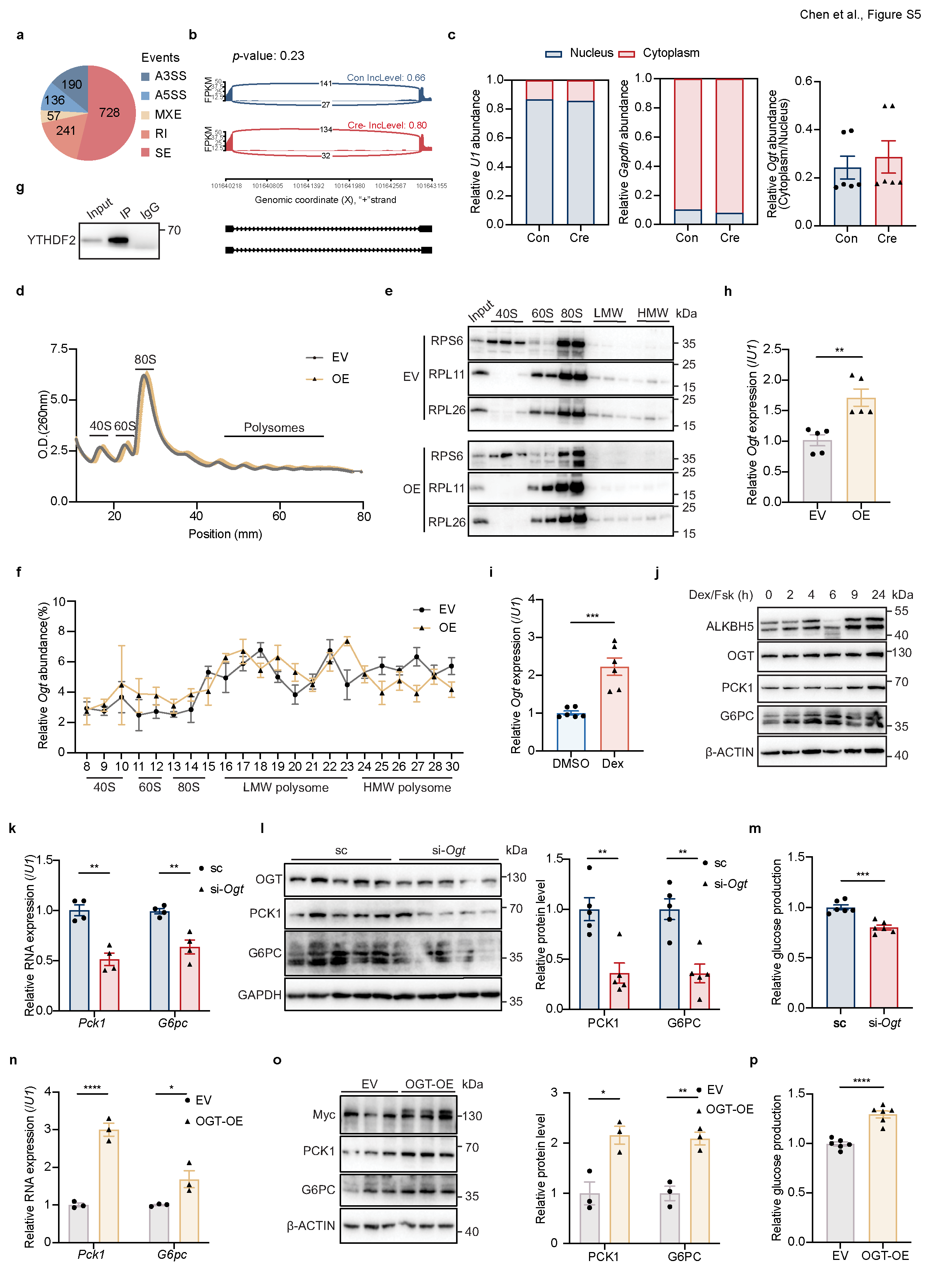

**Fig. S5 m^6^A-mediated regulation of *Ogt* and OGT-dependent control of hepatic gluconeogenesis. Related to Fig. 5.**

**a** Pie charts showing changes in alternative splicing patterns between Con and Cre MPH. **b** Sashimi plots illustrating the impact of ALKBH5 knockout on *Ogt* transcript splicing. **c** RT-qPCR analysis showing the impact of ALKBH5 knockout on the subcellular localization of *Ogt* RNA in MPH. *U1* and *Gapdh* were included as nuclear and cytoplasmic markers, respectively (n = 6). **d** Sucrose gradient-based polysome profiling of AML12 cells with or without ALKBH5 overexpression. **e** Western blot analysis to detect RPS6, RPL11, and RPL26 expression in ribosomal subunits (40S, 60S, 80S, and polysomes) from AML12 cells. **f** Distribution of *Ogt* mRNAs across polysome fractions in AML12 cells, quantified by RT-qPCR and normalized to input RNAs (n = 3). **g** Representative Western blot showing the efficiency of YTHDF2 immunoprecipitation in MPH. **h** RT-qPCR analysis showing the impact of ALKBH5 overexpression on *Ogt* expression in MPH (n = 5). **i** RT-qPCR analysis of *Ogt* expression in MPH with or without Dex treatment (n = 6). **j** Representative Western blot showing the expression of ALKBH5, OGT, PCK1 and G6PC in AML12 cells following Dex/Fsk treatment for the indicated time. **k–m** RT-qPCR analysis of *Pck1, G6pc* expression (**k**, n = 4), Western blot (left) and quantification (right) of PCK1 and G6PC proteins (**l**, n = 5) and glucose output assay (**m**, n = 6) in MPH transfected with scramble siRNA or *Ogt* siRNA. (**n**–**p**) RT-qPCR analysis of *Pck1, G6pc* expression (**n**, n = 3), Western blot (left) with quantification (right) of PCK1 and G6PC proteins (**o**, n = 3) and glucose output assay (**p**, n = 6) in MPH transfected with EV or OGT-overexpressing plasmid. Data are represented as mean ± SEM. Statistical significance was calculated by two-tailed unpaired Student’s t-test (**c**, **h**, **i**, **m**, **p**), multiple unpaired Student’s t-tests with Holm-Sidak correction (**k**–**l**, **n**–**o**), or two-way ANOVA followed by Sidak’s multiple comparisons test (**f**). **p* < 0.05, ***p* < 0.01, and ****p* < 0.001.

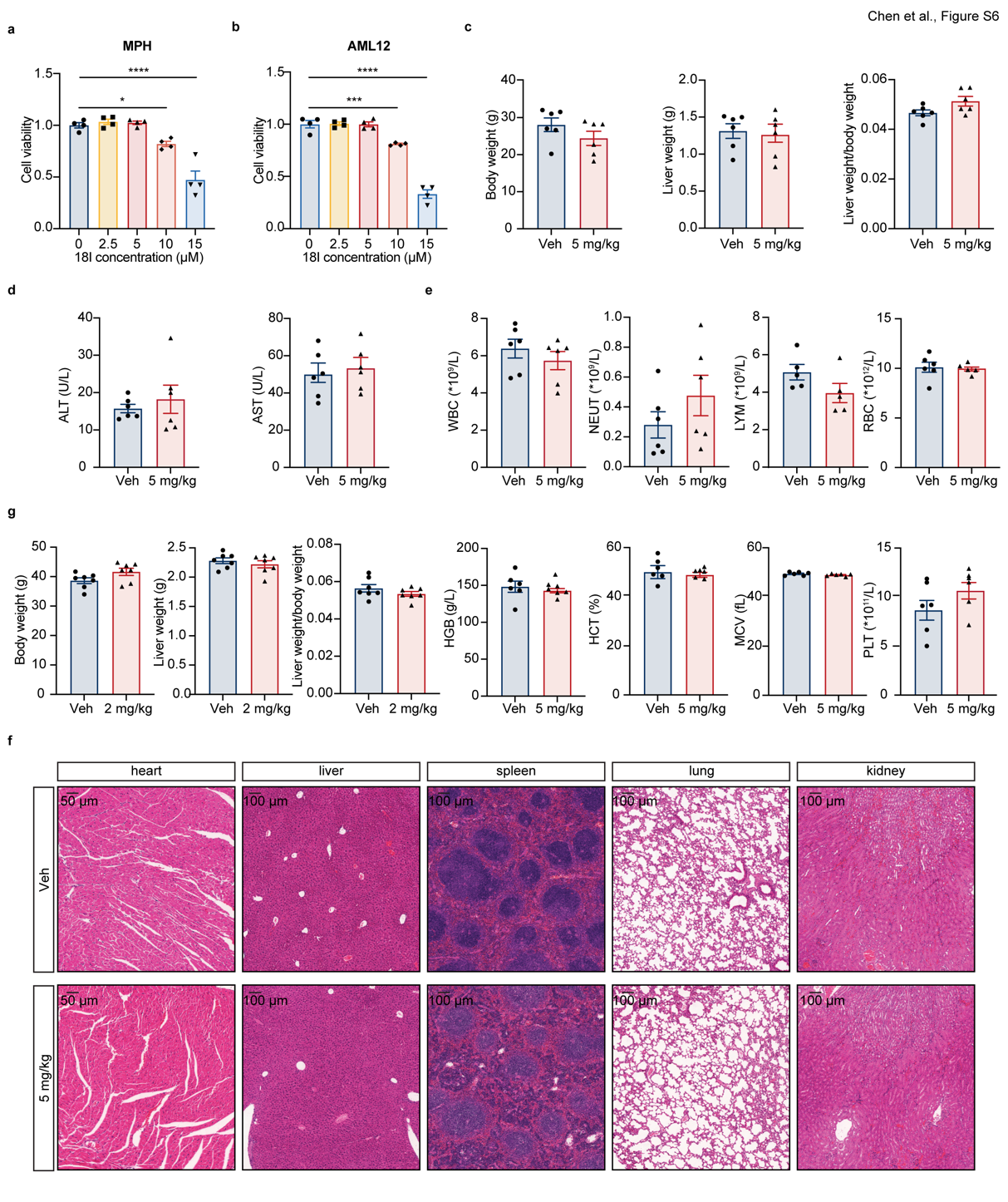

**Fig. S6 The ALKBH5 inhibitor 18l modulates gluconeogenesis and** **demonstrates *in vivo* safety in mice. Related to Fig. 6.**

**a** Cell viability of MPH treated with 18l at different concentrations (n = 4). **b** Cell viability of AML12 cells treated with 18l at different concentrations (n = 4). **c** Body weight (left), liver weight (middle), and liver weight-to-body weight ratio (right) in WT mice after injection of 18l for 14 days (n = 6). **d** ALT and AST levels measured in the plasma of WT mice after 18l injection (n = 6). **e** Complete blood count (CBC) of WT mice injected with 5 mg/kg 18l, showing standard hematological parameters including WBC, NEUT, LYM, RBC, HGB, HCT, MCV and PLT (n = 6). **f** Representative H&E staining of mouse heart, liver, spleen, lung, and kidney, showing tissue morphology in WT mice injected with 18l or vehicle (n = 3). Scale bar: heart: 50 μm, others: 100 μm. **g** Body weight (left), liver weight (middle), and liver weight-to-body weight ratio (right) in *db/db* mice after injection of 18l for 18 days (n = 7). Data are represented as mean ± SEM. Statistical significance was calculated by one-way ANOVA with Dunnett’s multiple comparisons test (**a**, **b**) and two-tailed unpaired Student’s t-test (**c**–**e**, **g**). **p* < 0.05, ***p* < 0.01, and ****p* < 0.001.
